# Burning down the mouse: Effects of wildfire and post-fire reseeding on Sin Nombre virus prevalence in its reservoir host

**DOI:** 10.64898/2026.09.02.748549

**Authors:** Carleen N. Silva, Frances M. Twohig, Matthew E. Gompper, Robert A. Nofchissey, Aaron C. Young, Steven B. Bradfute, Kathryn A. Hanley

## Abstract

Worldwide, physical habitats and biological communities are being reshaped by wildfire. Such changes have obvious potential to alter pathogen ecology, yet few studies, outside of those focused on ectoparasites, have tested the effects of wildfire on pathogen prevalence in wildlife. Even fewer have focused on the impacts of strategies for post-fire remediation on pathogen circulation. Here, we investigated the effect of wildfire and wildfire mitigation on the prevalence and viral load of Sin Nombre virus (SNV) infection in its reservoir host, the western deer mouse, using a study design of matched burned, unburned, and post-burn reseeded sites in northern New Mexico. In total, we screened 411 individual deer mice for SNV via RT-qPCR and analyzed the effect of wildfire and wildfire mitigation on relative mouse abundance, individual infection probability, site-level prevalence and viral load. Relative abundance was highest at reseeded sites, and model selection indicated that habitat structure – particularly the gradient from closed canopy to open-herbaceous habitat, was consistently associated with increased relative abundance. Similarly, SNV prevalence was significantly higher in reseeded sites than either burned or unburned sites but did not differ in burned versus unburned sites. Viral load did not differ between burn history types, and zero-inflated gamma hurdle models did not identify any strong predictors of viral load. Serendipitously, we were also able to investigate the impacts of an El Nino Southern Oscillation (ENSO) cycle on patterns of infection in a subsample of sites that were studied during and one year after an ENSO year and found that SNV prevalence increased significantly post-ENSO. Both reseeding and ENSO deliver resource pulses that can support increases in mouse density and thereby enhance SNV transmission. The impacts of post-fire management practices on SNV prevalence in its reservoir host that were revealed in this study should be considered when implementing such strategies and when utilizing treated areas.

**Open research statement:** To ensure transparency and reproducibility, the authors pledge to place all data and any novel code into Dryad upon acceptance of the manuscript.

## Introduction

Wildfires are increasing in both severity and frequency across the globe, driven by rising temperatures and shifting rainfall patterns (Cunningham et al., 2024; Dennison et al., 2014; Salguero et al., 2020; Westerling et al., 2006). Wildfires can dramatically reshape natural landscapes through reduction of ground litter, loss of canopy cover, and transformation of soil chemistry, among many other effects (Bansal, 2025). Following a wildfire, managers may implement strategies to prevent soil erosion, such as reseeding, replanting, mulching or chemical treatments. These interventions may alleviate some negative effects of wildfire, such as runoff or mudslides, but may exacerbate others, such as alteration of plant communities (Keeley, 2004). These wholesale changes in both burned and remediated landscapes have the potential to alter the transmission of pathogens among wildlife, thereby altering the risk of pathogen spillover to humans. In an increasingly flammable world, it is critical to gain an understanding of the effects of fire on wildlife infection.

Albery et al. (2021) have laid out multiple, intersecting mechanisms by which the consequences of wildfire could influence the exposure of wildlife to pathogens. At the nexus of this framework are the impacts of fire on host movement and resource availability. For directly-transmitted pathogens, wildfire may enhance inter-individual contact rates, and therefore transmission, via degradation of habitat complexity, concentration of existing resources, or production of resource pulses that support increases in the host population density. Alternatively, wildfire may reduce contact rates through habitat fragmentation, fire-mediated mortality, or production of resource pulses that disincentivize host foraging movements (Fay et al., 2025; Tonelli et al., 2026). Thus, wildfire may shape infection dynamics through multiple, potentially opposing, mechanisms. Although Albery et al. (2021) did not consider post-fire mitigation, it is clear that some management strategies may also affect host movement and host density. In particular, post-fire reseeding provides a large pulse in resources for granivores that may drive surges in their populations, and the plants that sprout from supplementary seeds may alter vegetation structure and plant community composition.

To date, the majority of studies investigating the impact of wildfire on the frequency of wildlife infection have focused on ectoparasites (reviewed in Scasta, 2015) or on proxies of disease risk, such as corticosteroid levels (Hing et al., 2017), immune function, or immune proteomes (Black et al., 2017; Bowen et al., 2015) rather than directly measuring wildfire impacts on pathogen prevalence. Two notable exceptions are work by Ecke et al. (2019) and Furtado et al. (2023). The former showed that wildfire-induced thermal stress may increase host susceptibility to parasitic infection in *Tropidurus oreadicus* lizards, and the latter demonstrated that reduced habitat complexity correlated with increased Puumala virus (PUUV, *Orthohantavirus puumalaense*) prevalence in bank voles.

In the current study, we investigated the effects of wildfires and post-fire management in northern New Mexico on Sin Nombre virus (SNV, *Orthohantavirus sinnombreense*) infection of western deer mice (*Peromyscus sonoriensis*). Western deer mice serve as a major reservoir host of SNV. Increases in SNV prevalence in deer mice are associated with increases in spillover of the virus to humans, in whom it can cause hantavirus cardiopulmonary syndrome (HCPS) with a fatality rate of ∼35% (Childs et al., 1994; Center for Disease Control, 2024). SNV circulation in deer mice is sensitive to ecological perturbations that alter mouse density, as the virus is transmitted via direct, aggressive contacts, primarily among adult males, that can increase with density (Mills et al., 1997; Pearce-Duvet et al., 2006). In particular, SNV prevalence in deer mouse populations, and subsequent human outbreaks, have been linked to El Niño Southern Oscillation (ENSO) events, which drive increased precipitation throughout the western United States, leading to mast seed production of piñon trees, a food resource pulse that supports increases in deer mouse density (Carver et al., 2015; Parmenter et al., 1993). While ENSO cycles provide a well-established example of how ecological perturbations can alter the SNV system within deer mice, wildfire and post-fire mitigation may act via similar mechanisms.

Populations of *Peromyscus* spp. have generally been shown to increase following both wild and prescribed fires, likely due to release of competition with members of their guild and increased foraging efficiency in the absence of ground litter (reviewed in Jones, 1992; Zwolak et al., 2010). Such increases in host abundance could accelerate SNV transmission and enhance prevalence. Consistent with this paradigm, Mull et al. (2023) reported that more rodents were seropositive for orthohantaviruses in burned prairies in the U.S. compared to clear cut or unmanaged control areas, and Gheler-Costa et al. (2022) found Brazilian Sigmodontinae rodents experienced much faster population growth and were more frequently positive for orthohantaviruses after sugarcane harvest fires than in areas where fire had been suppressed. However, fire effects are unlikely to be universal. Sharp Bowman et al. (2017) found that fires in the Great Basin, USA decreased *Peromyscus* population sizes, suggesting that severe fires may result in unsuitable habitat and decreased host abundance. Post-fire aerial reseeding may further modify these relationships. By providing an artificial resource pulse, reseeding may act in an analogous fashion to resource pulses associated with ENSO cycles, supporting higher densities of deer mice and elevating SNV prevalence (Figure 1). However, as reseeding also alters habitat composition and vegetation structure, its effects may not be mediated solely by host density.

**Figure 1.**
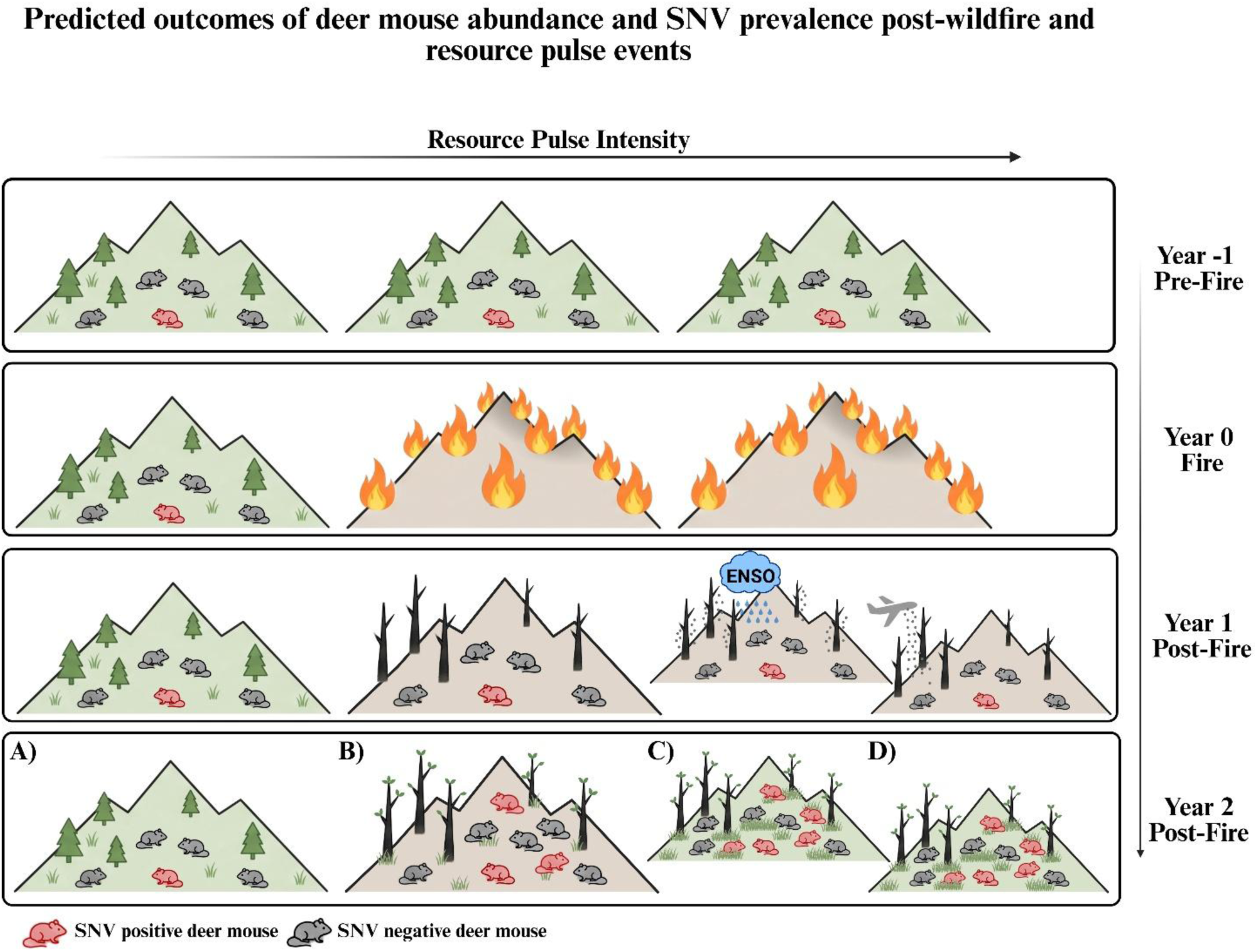
Predicted effects of wildfire and resource pulses due to reseeding or an ENSO event on vegetation, deer mouse abundance, and SNV prevalence in deer mice. The total number of mice represent the relative abundance of deer mice in a given burn history type; red mice represent SNV positive mice; black mice represent SNV negative mice. **A)** Independent of ENSO, we predict that unburned sites will remain unchanged in vegetation structure, deer mouse abundance, and SNV prevalence; **B)** Independent of ENSO, we predict that burned sites will experience substantial vegetation loss, increased mouse abundance and a concomitant increase in SNV prevalence due to wildfire; **C)** When ENSO occurs, we predict that burned sites will experience a pulse in seed production, which in turn will support even higher levels of deer mouse density than scenario B, and therefore higher SNV prevalence; **D)** When burned sites are subject to aerial reseeding, we predict that this resource pulse will act similarly to the natural resource pulse triggered by ENSO in scenario C.

These previous studies of deer mouse and SNV dynamics offer the rationale for the central hypothesis of this study, that changes in habitat structure and resource availability mediated by wildfire and post-fire management will lead to higher SNV prevalence within deer mouse populations via their effects on deer mouse abundance. Our study was conducted in northern New Mexico in the southwest United States, a region that has been ravaged by wildfires during the last decade. We tested several predictions stemming from our central hypothesis, namely: (1) relative abundance of deer mice and prevalence of SNV in deer mice would be higher in sites that had recently burned in a wildfire compared to ecologically-matched, unburned control sites (Figure 1A and 1B), and (2) relative abundance of deer mice and SNV prevalence would be higher in sites that had recently burned in a wildfire and been reseeded post-fire compared to burned sites that had not been reseeded and unburned sites (Figure 1D). Serendipitously, because 2023 was an El Niño year, we were also able to test (3) whether prevalence of SNV would increase at a one-year lag following an ENSO event (Figure 1C), as has been demonstrated previously in the desert southwest (Engelthaler et al., 1999; Yates et al., 2002). Additionally, we tested whether two other well recognized patterns in SNV prevalence also held true in our data: (4) SNV prevalence would be higher in adult male than female mice, and higher in adults than juveniles (Douglass et al., 2001; Mills et al., 1997), and (5) SNV prevalence would correlate with degree of wounding (Douglass et al., 2001). Finally, although we had no *a priori* reason to expect a difference, we nonetheless tested (6) whether SNV viral load in individual mice differed among burn history types. To address our predictions, we live-trapped deer mice over the summers in 2023 and 2024 at areas that had burned with high intensity one or two years previously and were or were not reseeded, as well as at ecologically-matched, unburned control sites. SNV infection of these mice was detected via RT-qPCR of collected lung tissues.

## Methods and Materials

### Ethical Approvals and Permits

All procedures on mice were approved by the New Mexico State University Institutional Animal Care and Use Committee (approval number 2404001125) and permission was given by the Jemez and Pecos Las Vegas Ranger districts to trap mice within their respective national forests. Field procedures were conducted following the Animal Care and Use guidelines of the American Society of Mammologists (Sikes and Gannon, 2011).

### Study Areas

This study was conducted in northern New Mexico (NM) (Figure 2A), an area that experiences mild summer temperatures with highs averaging 29°C, extreme winter temperatures dipping as low as -34°C, and seasonal precipitation occurring during summer monsoons and winter snowstorms (Figure 3). Northern NM has experienced multiple large fires in the last few years, including the two foci of this study, the Cerro Pelado fire and the Hermit’s Peak/Calf Canyon fire. The Cerro Pelado fire, which began in April 2022, seven miles east of the village of Jemez Springs, NM, USA (35° 46’ 30” latitude, -106° 35’ 4” longitude), was a mixed severity fire that burned over 45,000 acres (Figure 2B). The Hermit’s Peak fire started as a prescribed burn but was declared a wildfire on April 6, 2022. The Calf Canyon fire started as a pile burn holdover in January and re-emerged in April 2022. The two fires eventually merged into one, becoming the largest and most devastating fire in New Mexico state history. The Hermit’s Peak Calf Canyon complex fire burned over 341,000 acres in the Santa Fe National Forest and surrounding areas near Las Vegas, NM (105°20’19” latitude, 35°48’59” longitude; Figure 2D).

**Figure 2.**
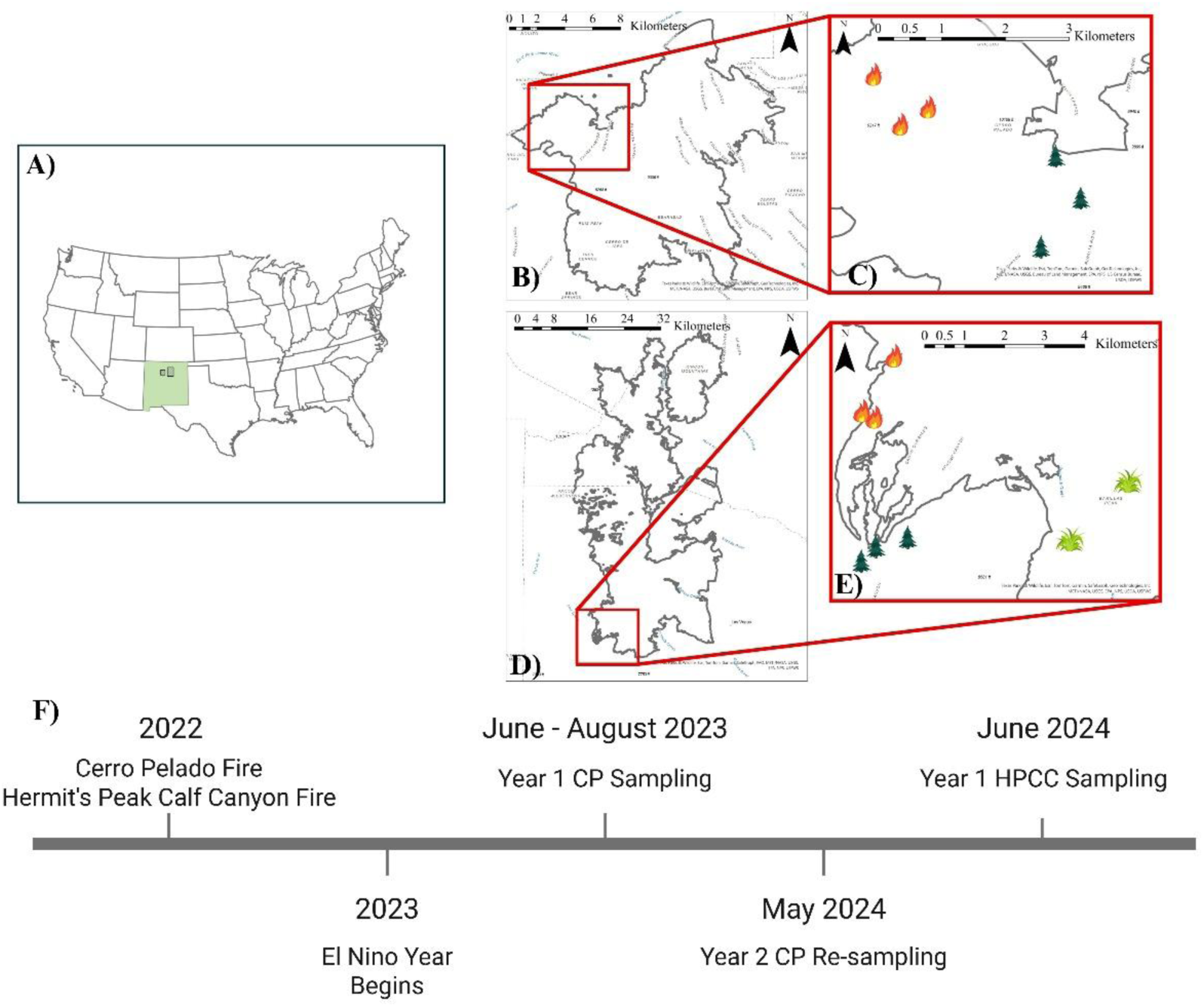
**A)** The state of New Mexico in the United States of America with locations of Cerro Pelado fire (left black box) and Hermit’s Peak Calf Canyon (right black box); **B)** Cerro Pelado fire perimeter; sampling area denoted by red box; **C)** Locations of burned and unburned sites. Burned sites represented by fire icons, unburned sites denoted by tree icons; **D)** Hermit’s Peak Calf Canyon fire perimeter; sampling area denoted by red box; **E)** Locations of burned, unburned, and reseeded sites. Burned sites represented by fire icons, unburned sites represented by tree icons, reseeded sites represented by grass icons; **F)** Sampling timeline; “CP” stands for Cerro Pelado; “HPCC” stands for Hermit’s Peak Calf Canyon. Map was created with ArcGIS Pro. Icons are from Canva free edition.

**Figure 3.**
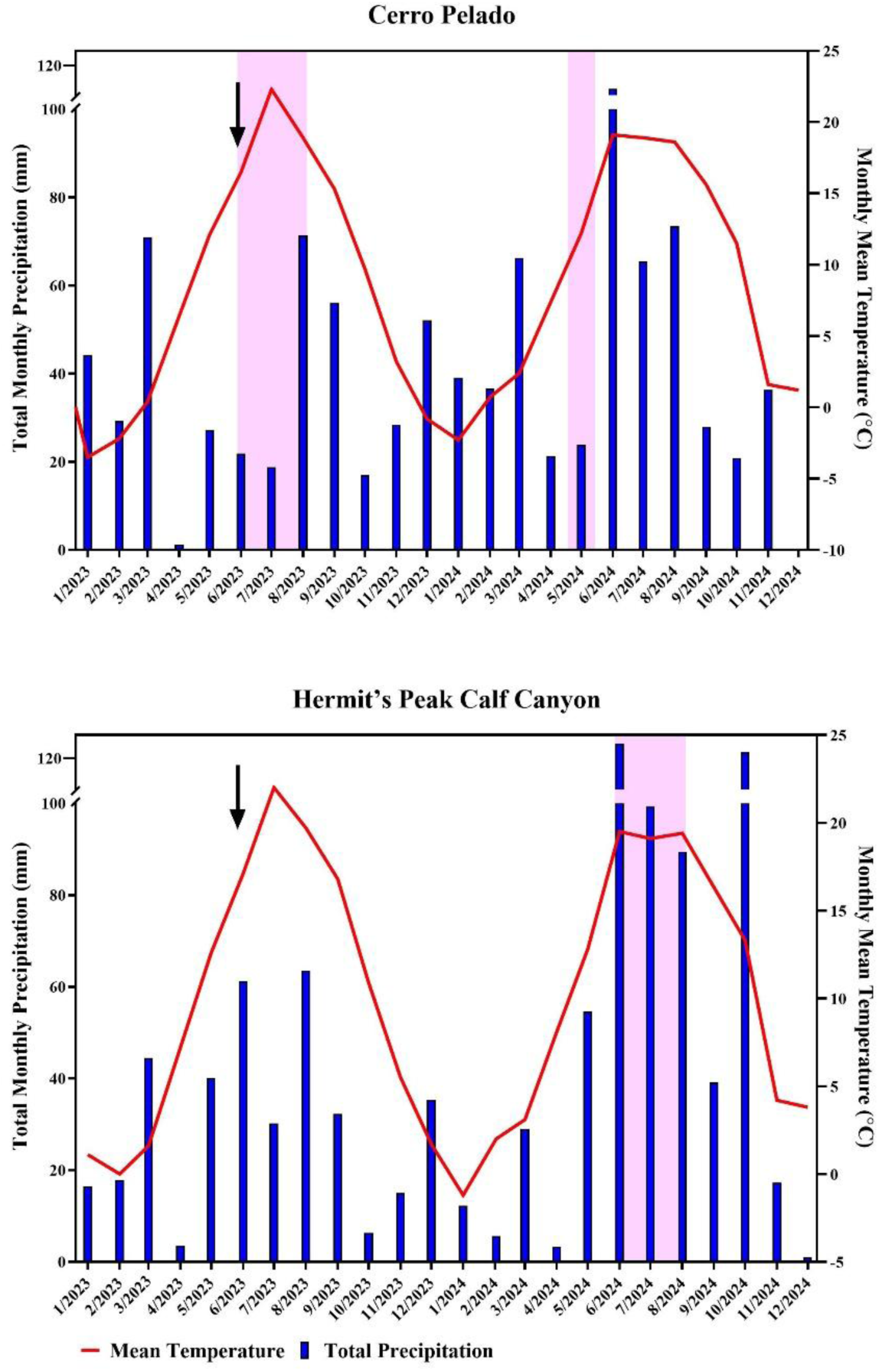
Climate data from January 2023 to December 2024 for **A)** Cerro Pelado and **B)** Hermit’s Peak Calf Canyon sampling areas. Sampling periods are indicated in pink shading. Incursion of El Niño is denoted with an arrow. Left y-axis (blue bars) shows total monthly precipitation. Right y-axis (red line) shows monthly mean temperatures. Climate data from Western Regional Climate Center (https://www.wrrc.dri.edu).

To assess the impact of habitat alterations due to fire and post-fire mitigation on relative mouse abundance and SNV prevalence and viral load, mice were live-trapped at 3 sites each in areas affected by the Cerro Pelado or the Hermit’s Peak-Calf Canyon fire, as well as 3 ecologically-matched, unburned control sites for each fire, and 2 sites that had burned and been reseeded, for a total of 14 sites (Figure 2C and 2E). Trapping was initially intended to occur at the Cerro Pelado fire sites in 2023 and the Hermit’s Peak-Calf Canyon fire sites in 2024 (Figure 2F). However, in 2023, the National Oceanic and Atmospheric Administration (NOAA) declared the arrival of an El Niño event during our sampling session (NOAA, 2023). To test the impact of El Niño on SNV prevalence in deer mice, the burned sites in the Cerro Pelado fire were resampled in 2024 (Figure 2F); time restrictions prevented the resampling of the unburned sites. Additionally, trapping was initially intended to occur at three reseeded sites, but extreme rainfall washed out the road to our final reseeded site, making it inaccessible. Instead, we re-sampled the site with the lowest trap success (unburned Site ‘U3’) to attempt to standardize representation across all burn history types. The data from the resampled site was not treated as an independent site, but as a continuation of the previous trapping iteration.

### Sampling Sites

Burned sites were constrained to areas categorized as high severity soil burn, determined by a post-fire BAER (burned action emergency response) survey conducted by the U.S. Forest Service (NM Fire Info 2022). To select burned sites, first 100 random points were generated within high severity burned areas that were within 500 meters of a road and spaced at least 250 meters apart. A buffer of 100 meters was created around the 100 points, and sites for which buffered areas were not completely within a high severity burn were excluded. To select unburned sites, 100 unburned points were randomly generated in areas that mirrored burned sites in terms of elevation and dominant vegetation, after which site selection followed the same protocol as for burned sites, save that unburned sites were also excluded if they experienced any burning after 1980, or if they had any visible burn scars upon visual inspection. Both burned and unburned sites that fit all criteria were assigned a number, and final sites were selected with a random number generator, with each burn type replicated three times, and separate control sites selected for each fire. Reseeded sites at Hermit’s Peak Calf Canyon were selected following the same protocol as for burned and unburned sites from an area that had been previously classified as an area of high severity burn and that had been aerially reseeded in 2022 (Figure 2E). The seed contained a mix of winter wheat (*Triticum aestivum*), cereal rye (*Secale cereale*), western wheatgrass (*Pascopyrun smithii*), and blue grama grass (*Bouteloua gracilis*) (National Resources Conservation Service, 2023).

### Animal collection and processing

A power analysis conducted prior to initiation of the study determined that the sample size necessary for a 70% power to detect a difference of 20% prevalence of SNV between burned and unburned sites, with sites within a burn category pooled, was 305 deer mice; thus every effort was made to achieve this sample size. Small mammals were trapped in Sherman Traps (H.B. Sherman Traps, Inc, Tallahassee, FL, USA) arranged in grids of 100 traps (10 lines with 10 traps, 10 meters apart) to allow estimation of relative mouse abundance. In 2023, to enhance the number of deer mice available for estimation of SNV prevalence, 40-50 traps were added in two trap lines within 300 meters of grids to each site. Individuals from these additional trap lines were recorded and screened for SNV but excluded from relative abundance analysis. Across all sampling periods, trapping effort totaled 5,319 trap nights, including 1859 trap nights in 2023, 853 trap nights at resampled sites in 2024, and 2,607 trap nights at Hermit’s Peak Calf Canyon sites in 2024. Traps were baited with peanut butter and oats wrapped in wax paper and furnished with 1-2 cotton balls for bedding. Traps were set in the morning (0500-0800 hrs) and ample shade was provided by piling vegetation on each trap. Traps were checked the following morning (0500-0800 hrs). Investigators followed trapping and sampling guidelines for trapping and sampling small mammals for virological testing provided by the CDC (Mills et al., 1995). Trapped deer mice were transported to a central processing area, where they were transferred into a plastic Ziploc bag and euthanized via chloroform inhalation. Other species (which were rarely encountered in this study) were released immediately from traps. Mice were assigned to an age class based on weight and coat pelage. Individuals that were >10 grams, were classified as juveniles or sub-adults. Individuals with grey pelage were identified as subadults, whereas individuals with brown pelage <10 grams were classified as adults. Sex was determined based on presence of internal sex organs. Reproductive status was identified based on presence of fetuses, distension of vagina, and lactation in females, and descended testes in males. Blood samples (0.01-1.0 mL) were collected via cardiac puncture into 1.5 mL eppendorf tubes and held on ice. Blood was centrifuged within 24 hours and serum was frozen on dry ice. Instruments were disinfected after processing each individual with REScue™ (Virox Technologies Inc., Oakville, ON, CAN) and 90% ethanol. The dissection field was also disinfected with REScue™ between individuals. Lung tissue was collected for analysis, placed in pre-labeled cryovials and immediately frozen on dry ice.

### RNA Extraction

Lung samples were transported on dry ice and stored at -80°C. RNA extractions were conducted in a biosafety cabinet in a BSL-2 laboratory with BSL-3 precautions. Samples were placed on dry ice while completing RNA extractions using Qiagen Viral RNA Kit (Qiagen, Hilden, GER), following manufacturer protocols with minor adjustments. Briefly, 20-40 mg of each lung sample was placed in bead beater tubes with 0.5 mg of 1.00 mm diameter zirconia beads (BioSpec, Bartlesville, OK, USA), 1.0 mg of 2.3 mm diameter zirconia beads (BioSpec, Bartlesville, OK, USA), and 900 mL of AVL buffer. Tissue was homogenized in a BioSpec MiniBeadBeater-19 (model 607) twice at 4,350 RPMs for 30 seconds, with a 60 second rest in between. Homogenates were centrifuged for seven minutes at 7000 RPMs, transferred to a 1.5 mL microcentrifuge tube and 5.6 µL of RNA carrier – re-suspended in AVE per manufacturer protocol – was added. Remaining RNA extraction steps were completed following manufacturer protocol, with RNA eluted in 40 µL of nuclease free water. RNA samples were stored at -80°C until cDNA synthesis.

### cDNA Synthesis

Synthesis was performed in a 1.5 mL microcentrifuge tube in a 20 µL reaction, composed of 5 µL of RNA template, 1 µL of random primers (ThermoFisher Scientific, Waltham, MA, USA), 1 µL of dNTP (10mM; ThermoFisher Scientific, Waltham, MA, USA), and 5 µL of RT-qPCR grade water (ThermoFisher Scientific, Waltham, MA, USA), which was incubated at 65°C for five minutes, placed on ice for two minutes, then centrifuged quickly to remove condensation. Then, 4 µL of 5x First Strand Buffer (ThermoFisher Scientific, Waltham, MA, USA), 2 µL of dithiothreitol (DTT) (0.1 mM; (ThermoFisher Scientific, Waltham, MA, USA), and 1 µL of RNAseOUT (ThermoFisher Scientific, Waltham, MA, USA) were added to each tube. Samples were incubated at 25°C for two minutes after which 1 µL of SuperScript II (ThermoFisher Scientific, Waltham, MA, USA) was added to each reaction. The reaction was incubated at 25°C for 10 minutes. Then, cDNA was synthesized at 42°C for 50 minutes. Synthesis was terminated at 70°C for 15 minutes. Lastly, cDNA samples were briefly centrifuged, labeled and stored at −20°C.

### RT-qPCR

RT-qPCR reactions were conducted using TaqMan™ Fast Advanced Master Mix (ThermoFisher Scientific, Waltham, MA, USA). Primers and probes (Integrated DNA Technologies, Inc., Coralville, IA, USA) used for the reactions can be found in Appendix S1:Table S1. Primers and probes were rehydrated in 0.1X Tris-ethylenediaminetetraacetic acid (TE) buffer to 100 mM. Each PCR plate contained a 10-fold standard dilution of 10^-1^ to 10^-8^ of cDNA from a Sin Nombre virus positive control. Additionally, plates contained no template controls (NTC). The probe was diluted to 1:10 and 1.5 µL was added to a 1.5 mL Eppendorf tube along with 3 µL of both primers and 22.5 µL of RT-qPCR water. This was diluted to 10X, then 180 µL of RT-qPCR water and 300 µL of TaqMan Fast Advanced Master Mix was added. 18 µL of the mixture was added to each well in a 96-well plate. 2 µL of sample cDNA, RT-PCR grade water (NTC), or known positive cDNA (positive control) was added to each well. All samples were run in duplicate and were separately screened for 18S ribosomal rRNA as an independent measure of sample RNA concentration with an endogenous control kit that included 18S primers and probes (ThermoFisher Scientific, Waltham, MA, USA). Conditions for RT-qPCR reactions were 95°C for 20 seconds, 95°C for 1 second, and 58°C for 30 seconds for 40 cycles, and samples with mean Ct values less than 40 were considered positive (Banther-McConnell et al., 2024). Samples that were ambiguous (one well positive and one well negative; 12% of total samples) were re-run until both wells were positive or negative in the same run. RT-qPCR results were analyzed on BioRad CFX Connect (Bio-Rad Laboratories, Hercules, CA). ΔCt was calculated as the difference between Ct values of SNV positive wells and 18S control wells.

### Habitat Data

To understand habitat variation within and among burn history types, five variables, which together accounted for 100% of ground cover, were used to characterize habitat: percent herbaceous ground cover, percent bare ground cover, percent tree canopy cover, percent shrub ground cover, and percent litter ground cover. Data for each were collected from a 200-meter radius around trap locations (distance chosen to account for spatial movements of deer mice; Douglass et al., 2006, Wood et al., 2010) through publicly available, remotely sensed data hosted through the Multi-Resolution Land Characteristics Consortium (MRLC) of the U.S. Geological Survey (Rigge et al., 2024). Since available data had a resolution of 30 pixels, the mean values were taken at 200 meters. Appendix S1:Table S2 shows the location, burn history type, and values for each of the six variables for each site in this study.

### Data Analysis

#### Habitat characterization

Due to the inherent correlation with ground cover categorization, all five habitat variables were centered and subject to a Principal Component Analysis (PCA) to reduce dimensionality. Burn history types were compared via an ANOVA on Principal Component 1 (PC1), and PC1 was used in generalized linear models, described below. PC1 was included in models to evaluate whether continuous variation in habitat structure was associated with variation in deer mouse abundance or SNV prevalence beyond categorical burn history type. Changes in PC1 in re-sampled areas were compared with a paired t-test, with sites paired across years.

#### Deer mouse relative abundance

Trap success in grids was used as a proxy for relative deer mouse abundance and was calculated for each site as the number of deer mice caught in a trapping grid per total number of trap nights (Mariën et al., 2024; Polito et al., 2022). Trap nights were defined as the difference between the number of traps set in grids and the number of traps falsely tripped during the session. Generalized linear models (GLM) were used to identify potential environmental and demographic variables associated with site-level abundance, excluding sites that had been resampled. Abundance was modeled with a beta regression, since abundance was treated as a proportion of successful traps between 0 and 1, with the fixed effects of (i) burn history type, (ii) year, (iii) proportion of trapped individuals identified as reproductive, and (iv) PC1, as well as the interactions of each. Year was included to account for interannual variation associated with post-fire recovery and climatic conditions, such as ENSO during the 2023 sampling period. An *a priori* model candidate list of 20 models (Appendix S1:Table S3) were run in R using package ‘betareg’ version 3.2-4 (Cribari-Neto & Zeileis, 2010). Akaike’s information criterion corrected for small sample sizes (AICc) was used to compare models. Models within two ΔAICc of each other, were considered similarly supported. Where appropriate, the most parsimonious model was selected to visualize effects across competitive models.

#### Sin Nombre Virus Infection

SNV infection was analyzed at two complementary scales: individual infection status and site-level prevalence. Individuals in the burned sites in the Cerro Pelado fire that were sampled in 2024 were excluded from these main analyses and only used to test for impacts of El Niño. Explanatory variables for all models were assessed for collinearity using Pearson’s correlation coefficient and excluded from use in the same model if the correlation value was > 0.7. Models within two ΔAICc of each other, were considered similarly supported. Where appropriate, the most parsimonious model was selected to visualize effects across competitive models.

To test the effect of burn category on probability of infection a binomial GLM with infection status as the response variable was used. GLM analysis was completed in R (R Core Team, 2025). Fixed effects included (i) sex, (ii) burn history type (burned, unburned or reseeded), (iii) body size corrected mass (residuals of the ratio of weight to hindfoot; Schulte-Hostedde et al., 2005), (iv) number of wounds, and (v) year, as well as their pairwise interactions. For GLM analysis, 26 *a priori* model candidates were tested (Appendix S1:Table S4).

Site-level prevalence was calculated as the proportion of individuals that tested positive at each site and modeled using a beta regression in R package ‘betareg’ as a generalized linear model (Cribari-Neto & Zeileis, 2010). Fixed effected included: (i) burn history type, (ii) deer mouse relative abundance, (iii) year, (iv) proportion of adult males at a site – calculated as the number of adult male deer mice relative to total number of deer mice at a site – and (v) PC1 from the habitat variable PCA along with their pairwise interactions. In total, 32 *a priori* model candidates were tested (Appendix S1:Table S5). However, since at two sites no individuals tested positive and beta regression models cannot accommodate zeros, a Smithson-Verkuilen (2006) adjustment for all prevalence data was performed to account for the zeros. 95% confidence intervals of effect sizes were calculated using R package ‘emmeans’ version 2.0.1 (Lenth & Piaskowski, 2026).

Viral load was assessed using a zero-inflated gamma hurdle model to predict ΔCt values. The hurdle model first estimated the probability of being infected as a binary outcome (1 if infected; 0 if not) and subsequently used a gamma model with logit link function to estimate the effect of covariates on viral load. Fixed effects included (i) burn history type, (ii) sex, (iii) body size corrected mass, (iv) the number of wounds, and (v) year, as a categorical variable. In total, 28 *a priori* model candidates were tested (Appendix S1:Table S6) in R package ‘glmmTMB’ version 1.1.14 (Brooks et al., 2017).

#### Effects of El Niño Southern Oscillation

Differences in relative abundance between ENSO conditions (Year 1; 2023) and post-ENSO (Year 2; 2024) were evaluated with a paired t-test, treating the site as the repeated measure, on natural log-transformed abundance. To assess the impact of ENSO on SNV in deer mice a contingency table analysis on pooled data from burned sites was used to compare overall prevalence during ENSO and post-ENSO. Within the post-ENSO year, prevalence was also compared between males and females, and between wounded and non-wounded individuals, using contingency table analyses.

#### Effects of Sex on SNV Infection

To assess the effects of sex on SNV prevalence in deer mice, univariate analyses were first completed on pooled prevalence among individuals across all years and sites. Then, sex was evaluated by burn history types across all years. Lastly, the effect of sex on SNV prevalence was assessed by year and fire separately.

#### Degree of Wounding

The association between SNV infection and degree of wounding in mice was assessed with a nominal logistic regression with the number of wounds as the independent variable and the binary infection status as the dependent variable.

All analyses were completed in R Version 4.1.2 (R Core Team, 2025). Data were visualized with GraphPad Prism version 11.0.0 for Windows (GraphPad Software, Boston MA, USA).

## Results

### Habitat Characterization

Principal component 1 (PC1) from the PCA of habitat variables accounted for ∼83% of the total variance, PC2 accounted for another 14.3%, and all remaining PCs explained <3% of the total variation (Table 1). Herbaceous ground cover and tree cover were largely responsible for the variation within PC1. Ecologically, PC1 represents the transition from closed canopy forests to open, herbaceous areas, with higher PC1 scores indicating lower canopy cover and greater herbaceous ground cover. Subsequent comparisons of sites with different burn histories utilized just PC1. At the Cerro Pelado fire, burned sites did not differ from unburned sites (ANOVA; F(1,4) = 4.89, p = 0.09). At the Hermit’s Peak Calf Canyon fire, sites in different burn history types differed significantly (ANOVA; F(2,5) = 40.60, p = 0.0008): unburned sites had significantly lower PC1 scores compared to both burned (p = 0.001) and reseeded sites (p = 0.002), but burned and reseeded sites did not differ from each other (p = 0.60).

**Table 1.**
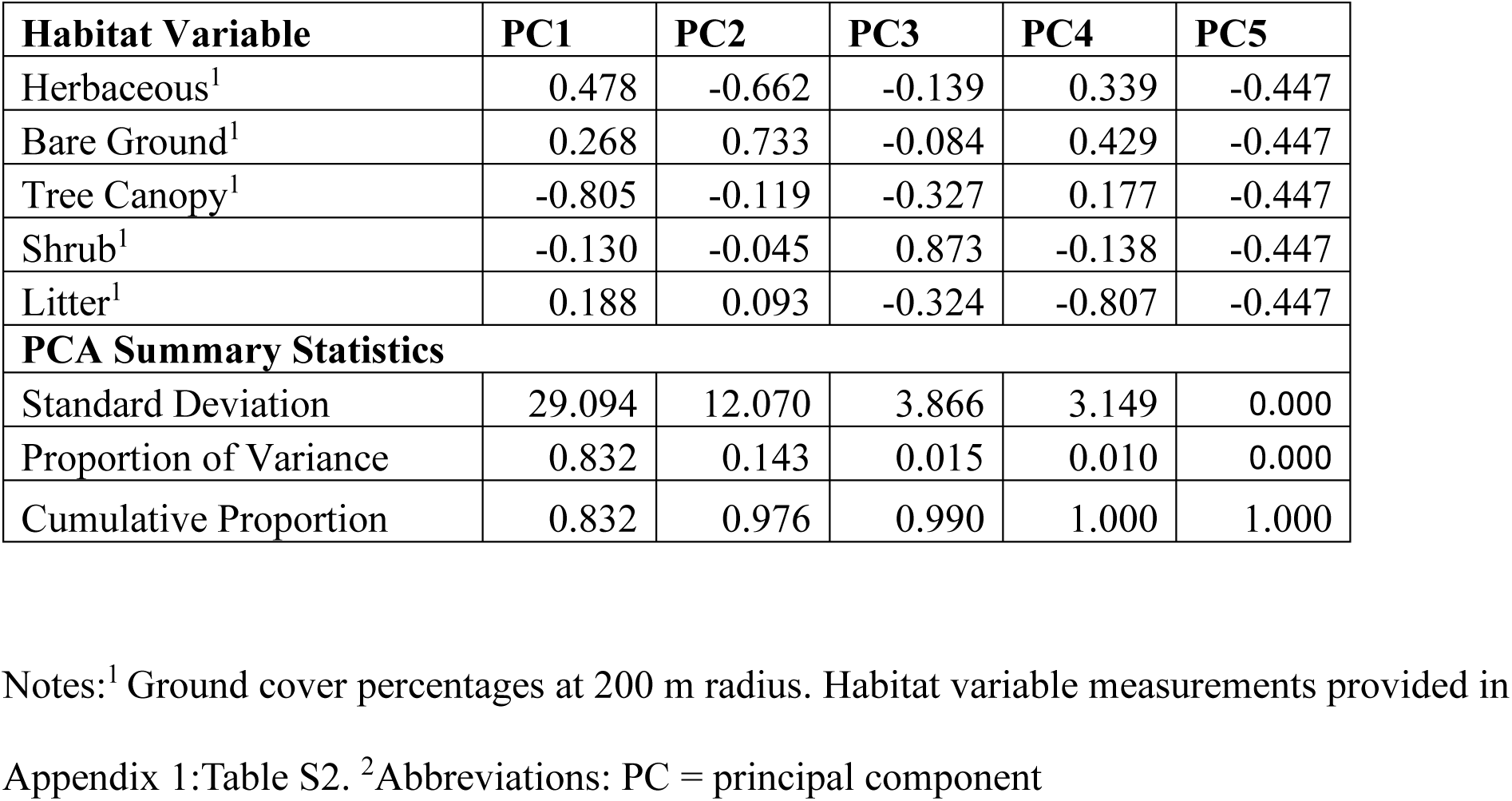
Principal component loadings summarizing major gradients of habitat structure on five habitat variables.

| Habitat Variable | PC1 | PC2 | PC3 | PC4 | PC5 |
| --- | --- | --- | --- | --- | --- |
| Herbaceous<sup>1</sup> | 0.478 | -0.662 | -0.139 | 0.339 | -0.447 |
| Bare Ground<sup>1</sup> | 0.268 | 0.733 | -0.084 | 0.429 | -0.447 |
| Tree Canopy<sup>1</sup> | -0.805 | -0.119 | -0.327 | 0.177 | -0.447 |
| Shrub<sup>1</sup> | -0.130 | -0.045 | 0.873 | -0.138 | -0.447 |
| Litter<sup>1</sup> | 0.188 | 0.093 | -0.324 | -0.807 | -0.447 |
| **PCA Summary Statistics** |  |  |  |  |  |
| Standard Deviation | 29.094 | 12.070 | 3.866 | 3.149 | 0.000 |
| Proportion of Variance | 0.832 | 0.143 | 0.015 | 0.010 | 0.000 |
| Cumulative Proportion | 0.832 | 0.976 | 0.990 | 1.000 | 1.000 |

Notes:<sup>1</sup> Ground cover percentages at 200 m radius. Habitat variable measurements provided in Appendix 1:Table S2. <sup>2</sup>Abbreviations: PC = principal component

PC1 scores did not differ significantly between years at the three burned sites that were sampled in both 2023 and 2024 (paired t-test; t(2) = 0.28, p = 0.80), although statistical power is low due to the small sample size. However, when considering herbaceous ground cover alone, mean herbaceous ground cover was greater in 2024 than in 2023 (paired t-test; t(2) = 4.45, p = 0.02) at resampled burned sites, as would be expected in sites recovering from a fire.

### Deer Mouse Relative Abundance

A total of 352 mice were caught in trapping grids across all years and all sites. Relative abundance differed among burn history types, with the highest observed relative abundance at reseeded sites (Figure 5A). A set of 20 GLM model candidates utilizing four explanatory variables to identify factors related to mouse relative abundance were evaluated through AICc (Appendix S1:Table S7). Three models fell within two ΔAICc of the highest ranked model (Table 2). Across all models, PC1 was the only shared predictor of relative abundance and was consistently positively associated with increased abundance (Estimate = 0.011 ± 0.004 SE, z = 2.876, p = 0.004). indicating higher mouse abundance in more open, herbaceous habitats (Figure 4). For visualization of the consistent relationship between PC1 and relative abundance, predictions from the PC1 only model indicated that relative abundance increased from 2.9 mice per 100 trap nights in the least open habitats, to 8 mice per 100 trap nights in the most open, herbaceous habitats, corresponding to a 2.8-fold increase. However, equally competitive models contained PC1 along with either (i) proportion of reproductive individuals at a site (Estimate = -1.8 ± 0.73, z = -2.47, p = 0.01) or (ii) occurring during the second year of sampling at the Hermit’s Peak Calf Canyon fire, (Estimate = 0.45 ± 0.17 SE, z = 2.62, p = 0.009), both of which additional effects were positively associated with greater mouse abundance.

**Figure 4.**
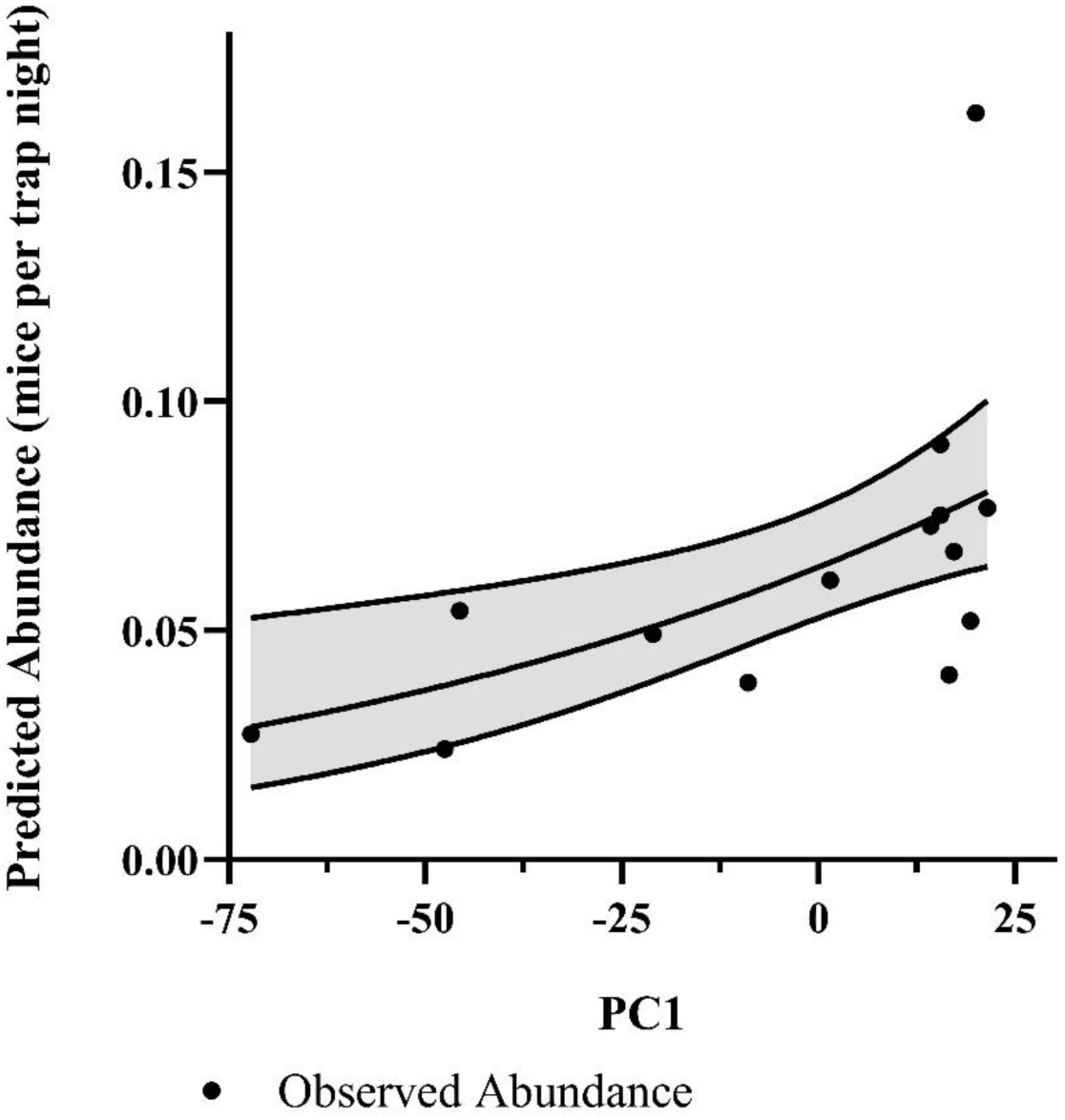
Predicted effect of PC1 on deer mouse abundance from beta regression analyses. Solid line represents model predictions, shaded regions indicated 95% confidence intervals, and points represent observed abundance values.

**Figure 5.**
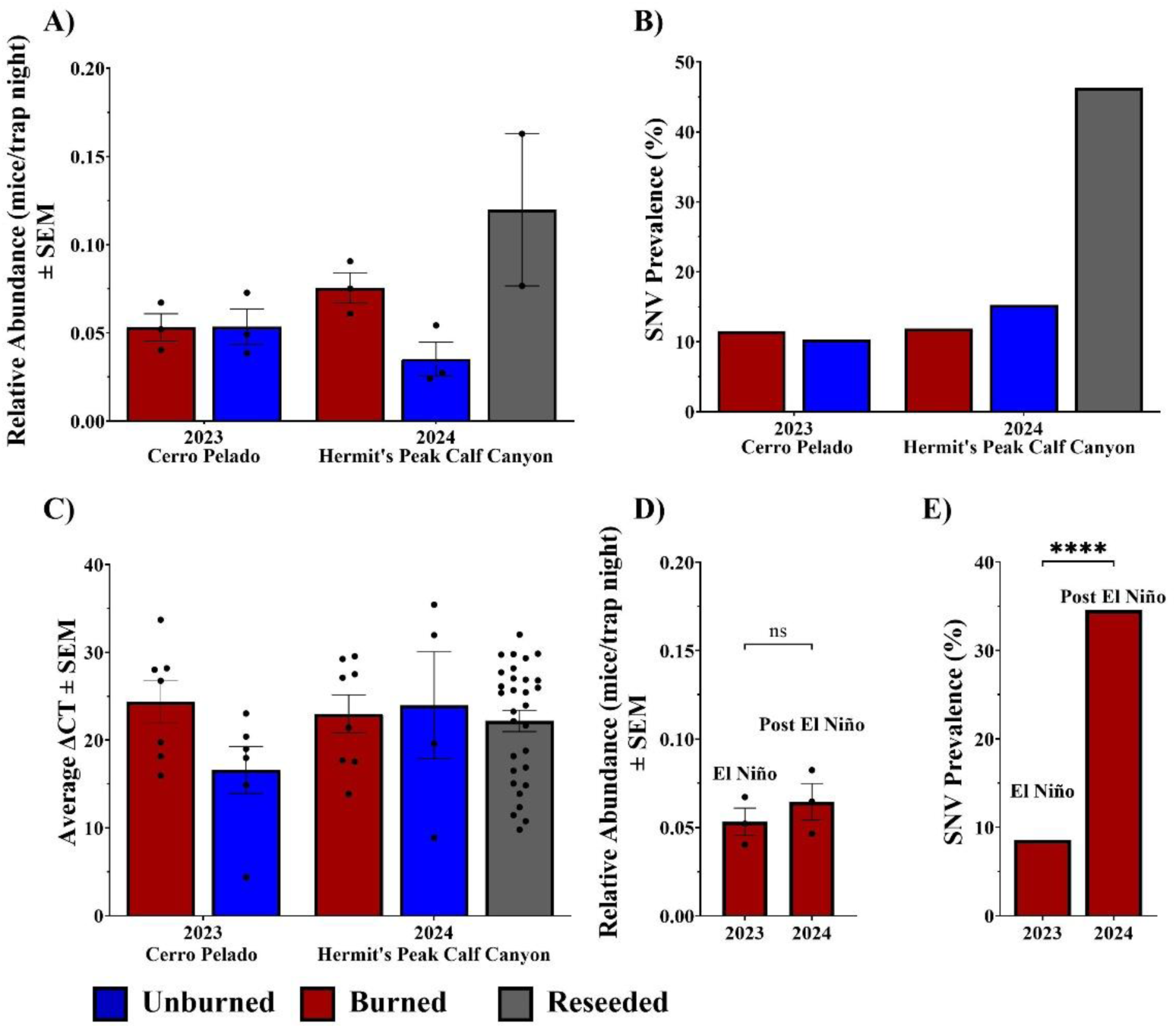
Wildfire effects on deer mice population and SNV dynamics. **A)** Mean relative abundance of deer mice across burn history types in 2023 at Cerro Pelado fire and in 2024 at Hermit’s Peak Calf Canyon fire; **B)** Observed pooled prevalence across burn history types in 2023 at the Cerro Pelado fire and 2024 at the Hermit’s Peak Calf Canyon fire; **C)** Mean viral load, quantified as ΔCt values, of SNV positive animals across burn history types in 2023 and 2024; **D)** Mean relative abundance of deer mice at burned sites in 2023 and 2024 at Cerro Pelado fire; **E)** Observed pooled prevalence for burned sites sampled during and one year after the 2023 El Niño event at the Cerro Pelado fire site. Significance bars represent results from paired t-test. Abbreviations: SEM = standard error of the mean; Ct = cycle threshold; ns = not significant; **** = p < 0.0001.

**Table 2.**
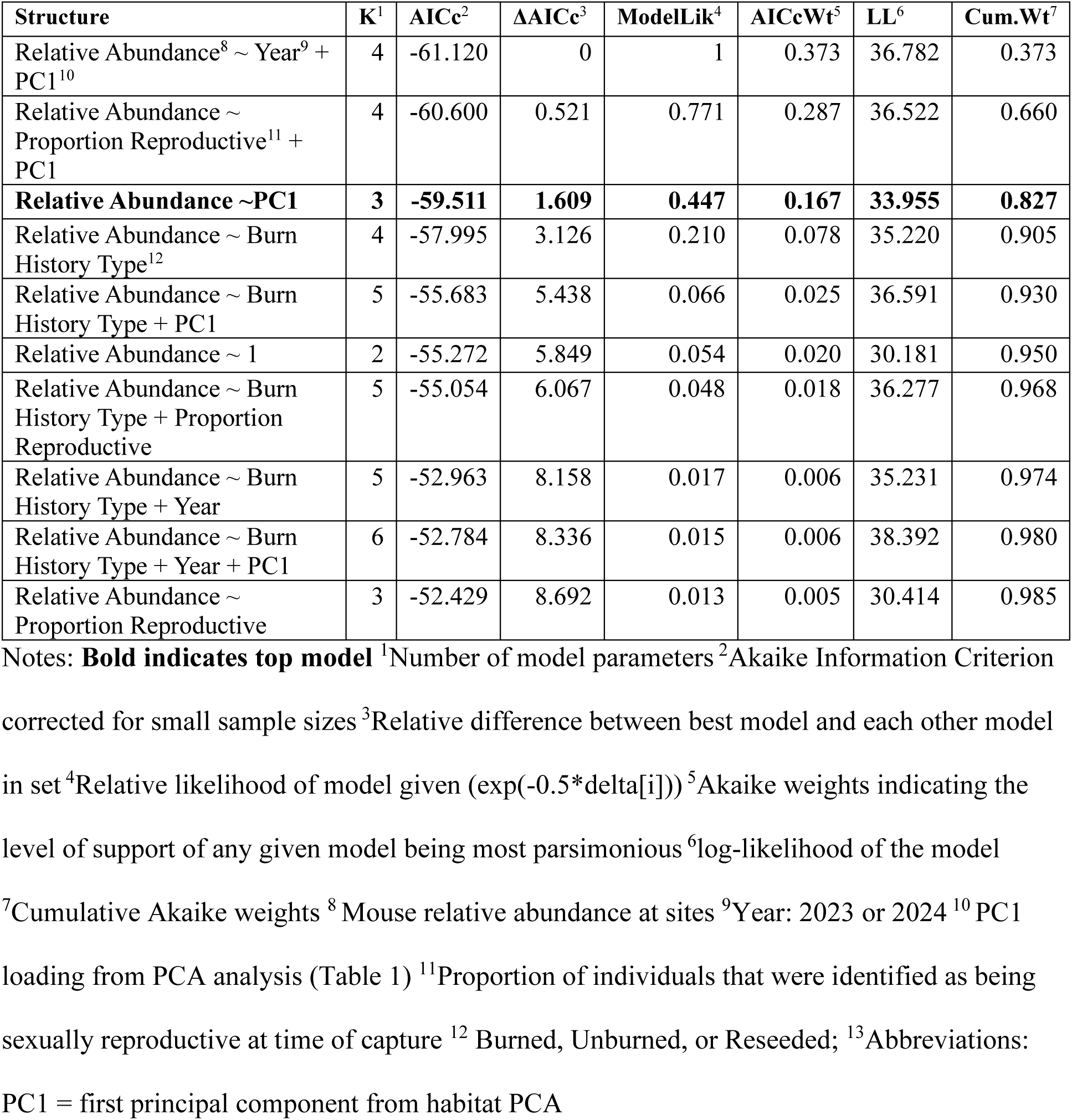
Top performing models predicting deer mouse abundance, with model selection indicating the strongest support for models including year, habitat structure (PC1), and proportion of reproductive individuals.

**Table 3.**
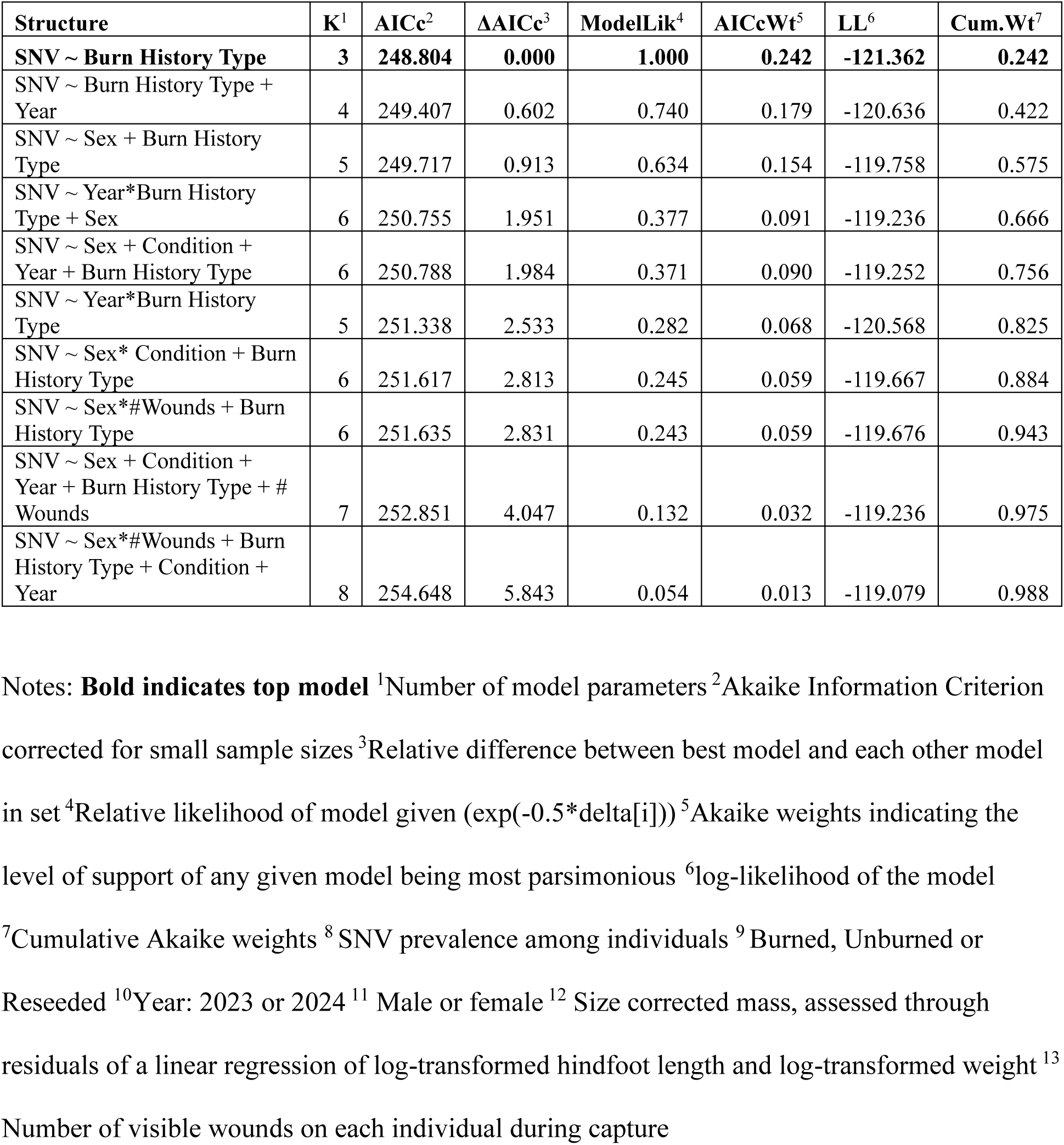
Top performing models predicting SNV infection among individuals, model selection indicating strongest support for models including burn history type, year, sex, and body condition.

### Sin Nombre Virus Prevalence Among Individuals

In addition to the 352 mice captured in grids, an additional 59 mice were captured in traplines outside of grids, and so we screened a total of 411 deer mice for SNV. Overall SNV prevalence at all Cerro Pelado sites pooled was 7.7%, while overall prevalence at Hermit’s Peak Calf Canyon sites was 24.6% (Figure 5B). The vast majority of infections (72/79) were detected in adults and therefore all subsequent analyses were limited to data from adult deer mice (n = 356).

In GLM analysis the most parsimonious, model within two ΔAICc included burn history type alone (Effect = 2.21 ± 0.36, z = 6.22, p = 5.14e-10). Deer mice at reseeded sites had 6.7 times higher odds of SNV infection relative to unburned sites (OR = 6.72, 95% CI: 3.04-15.99). Deer mice at burned sites did not have significantly higher odds of SNV infection than deer mice at unburned sites (OR = 0.73, 95% CI: 0.31-1.77). Although four other models fell within two ΔAICc all models consistently contained burn history type as a predictor (Appendix S1:Table S8). Sex, year, and size corrected mass appeared in the other competitive models but had less explanatory power than burn history type alone, and none were significant in their respective models.

### Sin Nombre Virus Prevalence Among Sites

Beta regression models identified a top ranked model that included only burn history type as an effect on SNV prevalence among sites (Table 4), with no additional models falling within two ΔAICc (Appendix 1:Table S9). The model predicted mean site-level SNV prevalence of 15% in unburned sites, 12.2% in burned sites, and 42.1% in reseeded sites (Figure 6). When effects of burn history type on SNV prevalence among sites were benchmarked against unburned sites as a control group in Table 5, burned sites did not differ from unburned sites (β = -0.24 ±0.31 SE, z = -0.78, p = 0.43), but prevalence at reseeded sites was significantly higher than either of the other two burn history types (β =1.42 ± 0.35 SE, z = 4.10, p < 0.0001). Burn history type accounted for a large proportion of variation in site-level prevalence (pseudo-R^2^ = 0.49).

**Figure 6.**
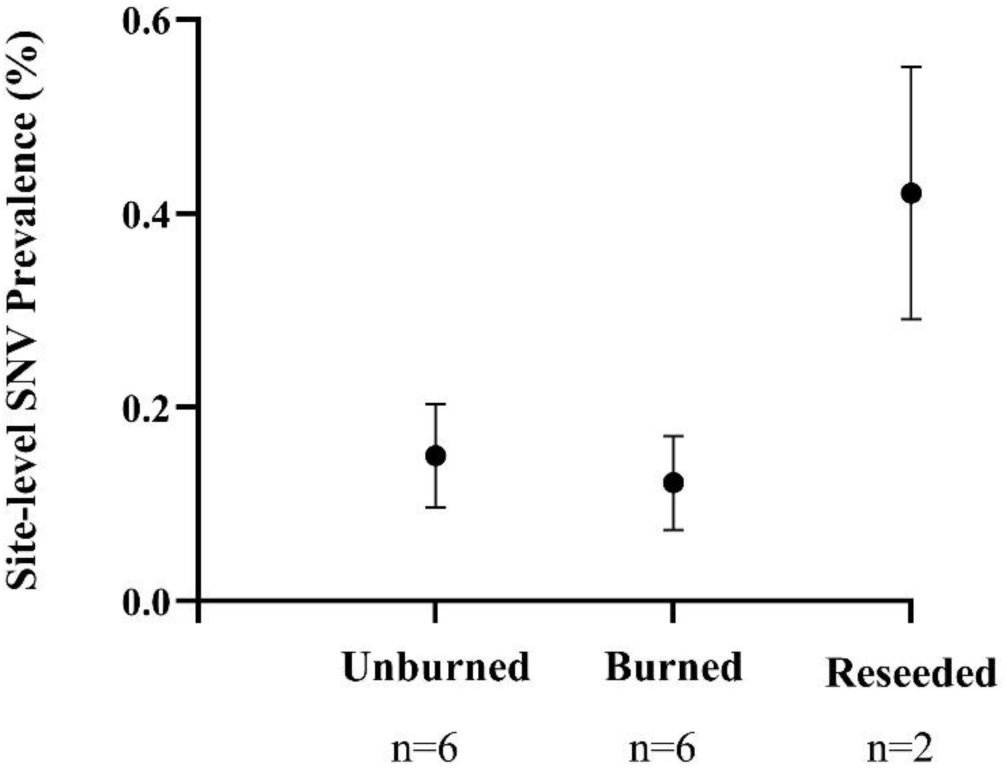
Predicted site-level SNV prevalence between burn history types, across both fires, from beta regression models. Points represent model estimates and error bars indicate 95% confidence intervals. Sample sizes beneath groups represent number of sites in each burn history type. Reseeded sites exhibited significantly higher SNV prevalence than either unburned or burned sites. Burned and unburned sites did not differ from one another.

**Table 4.**
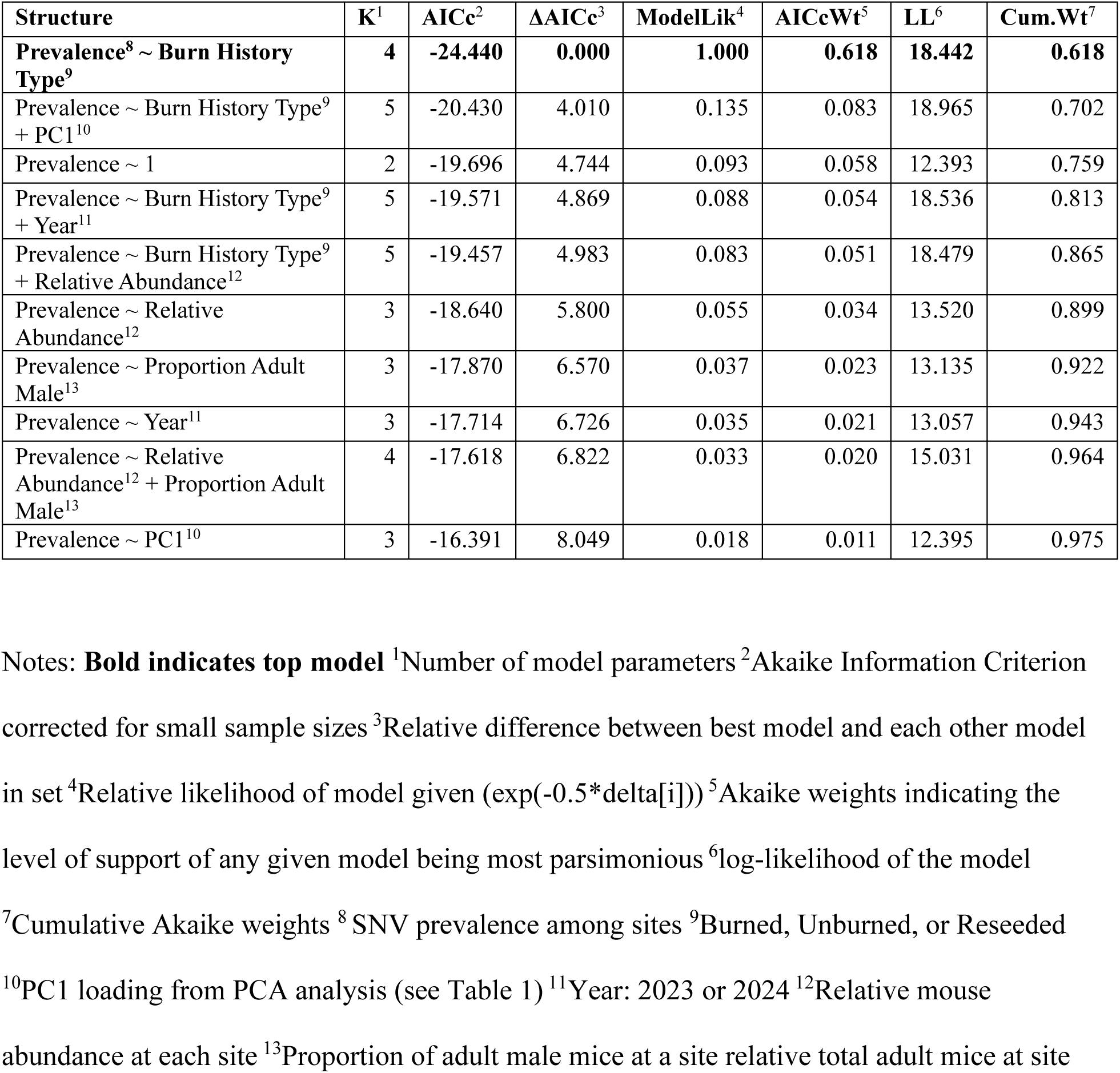
Top performing models predicting SNV prevalence among sites through a beta regression model with model selection indicating strongest support for model with burn history type as a fixed effect.

| Structure | K<sup>1</sup> | AICc<sup>2</sup> | ΔAICc<sup>3</sup> | ModelLik<sup>4</sup> | AICcWt<sup>5</sup> | LL<sup>6</sup> | Cum.Wt<sup>7</sup> |
| --- | --- | --- | --- | --- | --- | --- | --- |
| **Prevalence<sup>8</sup> ~ Burn History Type<sup>9</sup>** | **4** | **-24.440** | **0.000** | **1.000** | **0.618** | **18.442** | **0.618** |
| Prevalence ~ Burn History Type<sup>9</sup> + PC1<sup>10</sup> | 5 | -20.430 | 4.010 | 0.135 | 0.083 | 18.965 | 0.702 |
| Prevalence ~ 1 | 2 | -19.696 | 4.744 | 0.093 | 0.058 | 12.393 | 0.759 |
| Prevalence ~ Burn History Type<sup>9</sup> + Year<sup>11</sup> | 5 | -19.571 | 4.869 | 0.088 | 0.054 | 18.536 | 0.813 |
| Prevalence ~ Burn History Type<sup>9</sup> + Relative Abundance<sup>12</sup> | 5 | -19.457 | 4.983 | 0.083 | 0.051 | 18.479 | 0.865 |
| Prevalence ~ Relative Abundance<sup>12</sup> | 3 | -18.640 | 5.800 | 0.055 | 0.034 | 13.520 | 0.899 |
| Prevalence ~ Proportion Adult Male<sup>13</sup> | 3 | -17.870 | 6.570 | 0.037 | 0.023 | 13.135 | 0.922 |
| Prevalence ~ Year<sup>11</sup> | 3 | -17.714 | 6.726 | 0.035 | 0.021 | 13.057 | 0.943 |
| Prevalence ~ Relative Abundance<sup>12</sup> + Proportion Adult Male<sup>13</sup> | 4 | -17.618 | 6.822 | 0.033 | 0.020 | 15.031 | 0.964 |
| Prevalence ~ PC1<sup>10</sup> | 3 | -16.391 | 8.049 | 0.018 | 0.011 | 12.395 | 0.975 |

Notes: **Bold indicates top model** <sup>1</sup>Number of model parameters <sup>2</sup>Akaike Information Criterion corrected for small sample sizes <sup>3</sup>Relative difference between best model and each other model in set <sup>4</sup>Relative likelihood of model given ( $\exp(-0.5 \cdot \Delta AICc[i])$ ) <sup>5</sup>Akaike weights indicating the level of support of any given model being most parsimonious <sup>6</sup>log-likelihood of the model <sup>7</sup>Cumulative Akaike weights <sup>8</sup>SNV prevalence among sites <sup>9</sup>Burned, Unburned, or Reseeded <sup>10</sup>PC1 loading from PCA analysis (see Table 1) <sup>11</sup>Year: 2023 or 2024 <sup>12</sup>Relative mouse abundance at each site <sup>13</sup>Proportion of adult male mice at a site relative total adult mice at site

**Table 5.** Reseeded sites have significantly higher SNV prevalence, as demonstrated by the top performing model predicting SNV prevalence among sites.

| Fixed Effect | Estimate<sup>1</sup> | Standard Error | z-value | p-value |
| --- | --- | --- | --- | --- |
| Intercept | -1.73 | 0.21 | -8.15 | 3.78e-16 |
| Burned vs. Unburned | -0.24 | 0.31 | -0.78 | 0.43 |
| Reseeded vs. Unburned | 1.42 | 0.35 | 4.10 | 4.11e-05 |

Notes: Burned and reseeded sites are compared to unburned sites as a control group. <sup>1</sup> Log-transformed estimates.

### Individual and Site Level viral load

No models predicting viral load through a zero-inflated gamma hurdle model fell within two ΔAICc of the null model (Appendix1:Table S10). Roughly 83% of the data were structural zeros and filtered out when analyzing ΔCt values, therefore sample size was limited (n = 69). Viral load among SNV positive mice is shown in Figure 5C.

### Comparison of SNV prevalence during and one year after El Niño

In the resampled, burned sites of the Cerro Pelado fire, 53 deer mice were captured in 2024. Relative abundance did not differ between years (paired t-test: t=0.71, df = 2, p = 0.55; Figure 5E). SNV prevalence increased significantly between 2023 (the year El Niño occurred) at 6.93% and 2024 (one year post-El Niño) at 35.9%, (χ^2^ = 19.8, df = 1, p < 0.0001; Figure 5D). There were no differences between adult male and adult female prevalence in either pre-(χ^2^ = 2.92, df = 1, p = 0.09) or post-El Niño years (χ^2^ = 0.32, df = 1, p = 0.57).

### Comparison of SNV prevalence between sexes

Across all years and all sites, there was no difference in prevalence between adult males and adult females (χ^2^ = 0.28, df = 1, p = 0.60). However, at unburned sites, across all years, prevalence was significantly higher in adult males than adult females (χ^2^ = 5.35, df = 1, p = 0.03), while SNV prevalence did not differ between adult males and adult females in burned (χ^2^ =0.29, df = 1, p = 0.59) or reseeded sites (χ^2^ = 0.27, df = 1, p = 0.64).

### Comparison of SNV prevalence with degree of wounding

The number of wounds per mouse was not associated with SNV infection status (β = 0.12 ±0.16 SE, p = 0.44) and the regression explained little variation in SNV infection status (R^2^ = 0.001).

## Discussion

Wildfires in the southwestern USA are reshaping vegetation cover and resources availability, which may in turn alter infectious disease dynamics in wildlife. In the current study, we tested the hypothesis that wildfire effects on land cover would drive an increase in the abundance of deer mice and, as a consequence, an increase in the prevalence of SNV, for which deer mice are a major reservoir host. Additionally, we tested the hypothesis that re-seeding burned areas for post-fire mitigation strategy would result in even higher abundance of granivorous deer mice and, consequently, SNV prevalence, than would be found on burned but untreated land.

Serendipitously, we were also able to test whether deer mouse abundance and SNV prevalence increased on burned land after an El Niño event, a well-documented phenomenon in unburned, xeric habitats that has been attributed to the surge in piñon nut production supported by the uptick in rainfall during El Niño.

Importantly, we were able to replicate our study across two years and the sites of two major fires in New Mexico: the Cerro Pelado and the Hermit’s Peak Calf Canyon fires. We first assessed variation in ground cover among ecologically-matched sites with three different burn histories: burned, unburned, or burned and reseeded. Although both burned and reseeded areas had significantly lower tree cover, categorical burn history classification did not effectively capture variation in vegetation characteristics. This indicates that burn history type alone may only be a coarse level descriptor of habitat types in our study area and cannot fully account for heterogeneity present in post-fire landscapes.

Next, we tested our prediction that deer mouse abundance would differ between burn history types. However, habitat features, rather than burn history type *per se*, were most strongly associated with relative deer mouse abundance, consistent with previous studies showing that habitat characteristics outweigh burn variables in shaping rodent responses (Ecke et al., 2019). Deer mouse abundance increased as habitats became more open and herbaceous, reflecting this species’ preference for open habitats that provide benefits such as increased sources of food and protection from predators (Connolly & Orrock, 2018). While we were only able to sample two reseeded sites, limiting our ability to conduct robust statistical analyses with respect to this burn history type, deer mouse abundance at reseeded sites was more than twice as high as abundance in the other two burn history types.

SNV is transmitted among rodents by direct contact, so based on our *a priori* expectations regarding deer mouse abundance in different fire history types, we predicted that SNV prevalence would be highest at burned and reseeded sites relative to other land covers. For granivorous deer mice, re-seeding represents a form of resource provisioning, and resource provisioning has been shown to have a wide range of effects on wildlife disease, including transmission enhancement (Beck et al., 2026). SNV prevalence was analyzed at two levels of aggregation (individual infection and site-level prevalence). Burn history type was retained in the top-ranked models of both SNV prevalence among individuals and among sites in GLMs. In line with our prediction, reseeded sites had a higher proportion of infected individuals than either burned or unburned sites, possibly because the resource pulse delivered by aerial reseeding supported greater mouse reproduction, similar to ENSO events in this region that produce bumper crops of piñon seeds. Alternatively, mice may have moved into reseeded areas, driving up abundance. In either case, greater abundance would enhance rates of contact and thereby SNV transmission among mice. Because ENSO-driven increases in deer mouse abundance and infection have been linked to HCPS outbreaks in humans, it is noteworthy that an actively implemented management practice may have a similar effect. Moreover, within the US, wildfire frequency is increasing most rapidly in the west, which already experiences the majority of HCPS burden. Further investigating these overlapping areas of risk will become increasingly important for public health officials and forest managers, particularly if reseeding proceeds apace with wildfires.

In contrast to re-seeded sites, neither relative mouse abundance nor SNV prevalence differed between burned and unburned sites, contra our initial prediction and several previous studies (Zwolak et al., 2010; Culhane et al., 2022). Interestingly, a similar study in the Jemez Mountain range of New Mexico also found that deer mouse populations in burned areas did not differ from unburned areas one-to-three years after a large wildfire in 2011 (Peyton et al., 2025). Like the rest of the American west, New Mexico has experienced “megadrought” since 2000; this drought is thought to be one of the proximal causes of the Cerro Pelado and Hermit’s Peak Calf Canyon fires (Natural Resources Conservation Service, 2025). Since drought can limit population growth of *Peromyscus* species (Kuenzi et al., 2007; Nelson, 1993; Reed et al., 2007), it is possible that the impacts of the megadrought obscured any additional effects of fire on these populations. Moreover, wildfire may influence host pathogen dynamics through multiple competing pathways, in ways that may offset one another. Changes in resource availability and habitat structure may increase host abundance and transmission opportunities, while severe wildfires and prolonged drought may suppress host populations and reduce contact rates. Consequently, these opposing mechanisms may produce little net effect of wildfire alone on SNV prevalence.

As documented in numerous previous studies in xeric habitats (reviewed in Prist et al., 2017), Sin Nombre virus prevalence in deer mice did respond strongly to ENSO conditions at burned sites, which were the only sites samples during- and post-ENSO. SNV prevalence increased significantly, by ∼30%, one-year post-ENSO (2024) relative to the ENSO year (2023).

Herbaceous ground cover also increased at the sites sampled from 2023 to 2024, consistent with the understanding the ENSO alters SNV dynamics through alteration of food availability, mouse density, and mouse aggressive interactions. However, while we observed the expected increase in SNV prevalence, we did not observe the expected increase in deer mouse abundance post-ENSO. This may be due to timing of our field collection in the post-ENSO year, which occurred in late spring, prior to the period when deer mice tend to reach their peak populations. As a consequence, we may have failed to capture the full scope of population dynamics post-ENSO. In addition to ENSO, hantavirus prevalence has been linked to other shifts in environmental conditions, such as differences in habitat types (Kuenzi et al., 2001; Lehmer et al., 2012; Root et al., 1999; Yahnke et al., 2001), landscape structure (Langlois et al., 2001; Suzán et al., 2008), habitat quality (Pearce-Duvet et al., 2006), and level of human disturbance (Mackelprang et al., 2001). The observed increase in SNV prevalence post-ENSO provides independent support for the resource-pulse hypothesis that explains increased prevalence in reseeded areas. Together, these findings suggest that sporadic, episodic increases in resource abundance, either naturally or anthropogenically derived, may play a greater role in shaping SNV transmission than wildfire alone.

We had no *a priori* expectation that burn history would impact viral load, as viral replication is shaped by a product of a complex interplay among the genome, condition, immunity, sex and age of individual animals (Bagamian et al., 2013; Hannah et al., 2008). Indeed, none of the habitat or individual characteristics that we measured explained viral load. Notably, SNV RNA levels in deer mice tissue have been found to be extremely variable in laboratory settings (Schountz et al., 2012), which may obscure subtle ecological drivers of viral burden. Viral load has been previously correlated with the likelihood of transmitting SNV to an uninfected host (Bagamian et al., 2012), and based on our findings, infected mice across all burn history types would be equally likely to transmit SNV to uninfected mice.

Behavioral and population ecology, independent of habitat, can play a part in disease transmission. Our findings align with previous literature (reviewed in Khalil et al., 2014) and our own prediction, as SNV infection was almost exclusively detected in adult mice. Previous studies have also found that SNV prevalence is generally higher in male than female mice, but in the current study males had higher prevalence than females only in unburned sites. Further, this trend was largely driven by patterns at a single fire – Hermit’s Peak Calf Cayon – where no female mice in unburned sites tested positive for SNV. As our sample size for unburned sites at Hermit’s Peak Calf Canyon was limited (n=29), further study is needed to substantiate this finding. When data were aggregated more generally, sex was not a significant predictor of SNV prevalence. This result contrasts with expectations based on current understanding of SNV transmission ecology, as males are thought to drive transmission through aggressive interactions and thus are expected to exhibit higher prevalence (Root et al., 1999). This lack of sex-bias in infection may reflect temporal or demographic dynamics unique to our study or reflect the critical importance of habitat to SNV infection patterns. Though more wounds would seem to indicate more aggressive interactions, we found no evidence that an increase in wound count was correlated with increased infection probability..

There are several caveats to our findings. First, based on our *a priori* power analysis, our sample size may not have been sufficiently large to detect differences in SNV between burned and unburned sites with high power. Nonetheless, we did detect robust differences in SNV prevalence between reseeded sites and the other two burn history types, as well as a substantial impact of ENSO on SNV prevalence in a relatively small (n = 53) sample of mice. Second, we were only able to sample two reseeded sites, which may not adequately capture variation in the reseeded landscapes. Third, long-term studies (>10 years) of SNV ecology show evidence of a lag between peak deer mouse population density and peak SNV prevalence – a phenomenon termed delayed density dependence (Douglass et al., 2001; Luis et al., 2015; Yates et al., 2002). Shorter studies, like ours, may fail to detect the relationship altogether, particularly at local scales (Carver et al., 2011). Finally, our sampling took place one- to two-years post-fire. Sampling over a longer time period may reveal additional nuances to the effects of fire and post-fire management on SNV prevalence.

This study did reveal that a post-fire mitigation strategy – reseeding – enhanced deer mouse abundance and SNV prevalence. The similarities in the responses to resource pulses generated by reseeding and by ENSO raise important questions about how post-fire management actions may alter disease transmission in ecologically sensitive host-pathogen systems, such as SNV. The lack of differences in SNV prevalence observed between burned and unburned areas may indicate that wildfire may be less important than sporadic resource pulses to reservoir abundance and disease transmission. Finally, our documentation of exceptionally high SNV prevalence in deer mice in reseeded areas offers a critical caution for land managers and others utilizing reseeded landscapes. Our findings suggest that future research on the impacts of wildfire on wildlife disease ecology should incorporate the effects of the various treatments utilized to stabilize soils and slow runoff in the aftermath of the fire.

## Acknowledgements

Funding for this work was provided through the NSF GRFP, which funded C. Silva, the Joint Fire Science Program Award no. 23-1-01-32. This work was supported, in part, by USDA National Institute of Food and Agriculture NextGen program, award no. 2023-70440-40158, and the USDA-McIntire-Stennis award. The American Museum of Natural History Theodore Roosevelt Memorial Grant provided support for field work. We acknowledge laboratory and field assistance from Josiah Martinez, Eliana Ortega, Kaleb Villegas, Celia Lopez, and Eva Rangel. We thank Drs. Robert Parmenter and Theresa Laverty for their gracious loan of Sherman traps, and Dr. Stephanie Seifert for donating SNV positive RNA. We would also like to thank Dr. Jeanne Faire at Los Alamos National Laboratories and Drs. Justine Garcia and Sarah Corey-Rivas at New Mexico Highlands University for their logistical support during field seasons.

## Author Contributions

C. N. S., M.E.G., and K.A.H. designed the field study, R.A.N. and S.B.B. designed molecular methodologies. C.N.S. conducted field work trapping deer mice. C.N.S., R.A.N. and S.B.B conducted RNA extractions from lung samples. F.M.T. designed the SNV primers and probes used in RT-qPCR methodology and trained C.N.S in techniques used. C.N.S. conducted all RT-qPCR assays. A.Y. contributed to statistical analysis. All authors contributed to and approved the final manuscript.

