## Appendix 1 for "Burning down the mouse: Effects of wildfire and post-fire reseeding on Sin Nombre virus prevalence in its reservoir host"

Journal Name: Ecosphere

Manuscript Type: Article

Subject Track: Disease Ecology

Manuscript Title:

Burning down the mouse: Effects of wildfire and post-fire reseeding on Sin Nombre virus prevalence in its reservoir host

Authors names for publication:

Carleen N. Silva[1,2], Frances M. Twohig[3], Matthew E. Gompper[2], Robert A. Nofchissey[3], Aaron C. Young[2], Steven B. Bradfute[3] and Kathryn A. Hanley[1*]

Appendix S1

Table S1. Primers and Probes used for RT-qPCR.

| **Primers and Probes** | **Sequence** | **Length (base pairs)** |
|---|---|---|
| SNV Forward Primer | GGATTGATGACTTCCTGGCTG | 21 |
| SNV Reverse Primer | CCTTTGACTCATCAACCTGTAA | 22 |
| SNV Probe (with FAM fluorophore) | ACCCTAGAGATGCTGCATTGGCAA | 24 |
| 18S rRNA endogenous Control (with FAM fluorophore) | Commercial assay kit with primers and probes (Applied Biosystems Cat No. 4333760T) | |

Table S2: Habitat characteristics and locations for all sites.

| Site ID | Year | Type[1] | Easting[2] | Northing[2] | % Herbaceous[3] | % Bare[3] | % Tree[3] | % Shrub[3] | % Litter[3] |
|---|---|---|---|---|---|---|---|---|---|
| H35 | 2023 | Burned | 358507 | 3961018 | 39.80 | 28.11 | 4.24 | 5.00 | 22.84 |
| H36 | 2023 | Burned | 358869 | 3961258 | 34.47 | 37.67 | 2.87 | 5.00 | 19.99 |
| H37 | 2023 | Burned | 357964 | 3961585 | 47.21 | 18.75 | 2.91 | 5.00 | 26.13 |
| U1 | 2023 | Unburned | 360873 | 3960560 | 43.05 | 1.01 | 39.20 | 10.41 | 6.33 |
| U2 | 2023 | Unburned | 360641 | 3959105 | 43.77 | 4.07 | 27.94 | 8.92 | 15.30 |
| U3 | 2023 | Unburned | 361040 | 3959879 | 64.92 | 2.20 | 11.53 | 6.02 | 15.34 |
| H35 | 2024 | Burned | 358507 | 3961018 | 52.05 | 14.89 | 5.85 | 7.00 | 20.21 |
| H36 | 2024 | Burned | 358869 | 3961258 | 56.18 | 13.61 | 5.53 | 7.00 | 17.69 |
| H37 | 2024 | Burned | 357964 | 3961585 | 58.26 | 8.45 | 5.51 | 7.00 | 20.78 |
| H4 | 2024 | Burned | 450832 | 3938583 | 50.30 | 3.96 | 18.07 | 12.92 | 14.75 |
| H5 | 2024 | Burned | 451458 | 3940087 | 46.97 | 19.99 | 6.05 | 7.31 | 19.68 |
| H6 | 2024 | Burned | 451458 | 3940087 | 46.97 | 19.99 | 6.05 | 7.31 | 19.68 |
| S1 | 2024 | Reseeded | 457329 | 3936889 | 44.11 | 32.85 | 0.39 | 6.95 | 15.70 |
| S2 | 2024 | Reseeded | 455862 | 3935486 | 52.13 | 19.35 | 3.10 | 6.97 | 18.45 |
| U4 | 2024 | Unburned | 451050 | 3935343 | 18.10 | 2.53 | 56.80 | 16.58 | 6.00 |
| U5 | 2024 | Unburned | 450722 | 3934879 | 3.27 | 0.57 | 78.95 | 10.95 | 6.25 |
| U6 | 2024 | Unburned | 451817 | 3935565 | 16.47 | 0.80 | 51.55 | 25.01 | 6.18 |

Notes:

[1] Burn history type
[2] Universal Transverse Mercator (UTM) coordinates
[3] Ground cover percentages within a 200-meter radius

Table S3. 20 *a priori* model candidates for generalized linear models predicting relative mouse abundance with a beta distribution.

| **Structure** |
|---|
| Relative Abundance ~ 1 |
| Relative Abundance ~ Burn History Type |
| Relative Abundance ~ Year |
| Relative Abundance ~ PC1 |
| Relative Abundance ~ Proportion Reproductive |
| Relative Abundance ~ Burn History Type + Year + PC1 + Proportion Reproductive |
| Relative Abundance ~ Burn History Type + Year + PC1 |
| Relative Abundance ~ Burn History Type + Year |
| Relative Abundance ~ Burn History Type + PC1 |
| Relative Abundance ~ Burn History Type + Proportion Reproductive |
| Relative Abundance ~ Proportion Reproductive + Year |
| Relative Abundance ~ Proportion Reproductive + PC1 |
| Relative Abundance ~ Year + PC1 |
| Relative Abundance ~ Burn History Type*Proportion Reproductive |
| Relative Abundance ~ Burn History Type*Proportion Reproductive + PC1 |
| Relative Abundance ~ Burn History Type*Proportion Reproductive + Year |
| Relative Abundance ~ Burn History Type*PC1 |
| Relative Abundance ~ Burn History Type*PC1 + Proportion Reproductive + Year |
| Relative Abundance ~ Burn History Type*PC1 + Proportion Reproductive |
| Relative Abundance ~ Burn History Type*PC1 + Year |

Abbreviations: PC1 = first principal component from PCA analysis

Table S4. 26 *a priori* model candidates predicting SNV prevalence among individuals

| **Structure** |
|---|
| SNV ~ 1 |
| SNV ~ Sex |
| SNV ~ Burn History Type |
| SNV ~ Condition |
| SNV ~ # Wounds |
| SNV ~ Year |
| SNV ~ Sex + Condition + Year + Burn History Type + # Wounds |
| SNV ~ Sex + Condition + Year + Burn History Type |
| SNV ~ Sex + Condition + Year |
| SNV ~ Sex + Condition |
| SNV ~ Sex + Year |
| SNV ~ Sex + Burn History Type |
| SNV ~ Sex + # Wounds |
| SNV ~ Burn History Type + Year |
| SNV ~ Sex*Condition |
| SNV ~ Sex*#Wounds |
| SNV ~ Sex*Condition + Burn History Type + # Wounds + Year |
| SNV ~ Sex*Condition + Year |
| SNV ~ Sex*Condition + Burn History Type |
| SNV ~ Sex*Condition + # Wounds |
| SNV ~ Sex*#Wounds + Burn History Type + Condition + Year |
| SNV ~ Sex*#Wounds + Burn History Type |
| SNV ~ Sex*#Wounds + Condition |
| SNV ~ Sex*#Wounds + Year |
| SNV ~ Year*Burn History Type |
| SNV ~ Year*Burn History Type + Sex |

Table S5. 32 *a priori* model candidates predicting SNV prevalence among sites

| **Structure** |
|---|
| Prevalence ~ 1 |
| Prevalence ~ Burn History Type |
| Prevalence ~ Mouse Abundance |
| Prevalence ~ Year |
| Prevalence ~ Adult Male Prevalence |
| Prevalence ~ PC1 |
| Prevalence ~ Burn History Type + Mouse Abundance + Year + Adult Male Prevalence + PC1 |
| Prevalence ~ Burn History Type + Mouse Abundance + Year + Adult Male Prevalence |
| Prevalence ~ Burn History Type + Mouse Abundance + Year |
| Prevalence ~ Burn History Type + Mouse Abundance |
| Prevalence ~ Burn History Type + Year |
| Prevalence ~ Burn History Type + PC1 |
| Prevalence ~ PC1 + Mouse Abundance |
| Prevalence ~ PC1 + Adult Male Prevalence |
| Prevalence ~ Mouse Abundance + Adult Male Prevalence |
| Prevalence ~ Mouse Abundance + Adult Male Prevalence + Year |
| Prevalence ~ PC1 + Mouse Abundance + Adult Male Prevalence + Year |
| Prevalence ~ Burn History Type*PC1 |
| Prevalence ~ Burn History Type*PC1 + Mouse Abundance + Year + Adult Male Prevalence |
| Prevalence ~ Burn History Type*PC1 + Mouse Abundance + Year |
| Prevalence ~ Burn History Type*PC1 + Year + Adult Male Prevalence |
| Prevalence ~ Burn History Type*PC1 + Mouse Abundance + Adult Male Prevalence |
| Prevalence ~ Burn History Type*PC1 + Mouse Abundance*Adult Male Prevalence |
| Prevalence ~ Burn History Type*Adult Male Prevalence |
| Prevalence ~ Burn History Type*Mouse Abundance |
| Prevalence ~ Adult Male Prevalence*Mouse Abundance |
| Prevalence ~ Adult Male Prevalence*Mouse Abundance + Year + PC1 + Burn History Type |
| Prevalence ~ Adult Male Prevalence*Mouse Abundance + Year + PC1 |
| Prevalence ~ Adult Male Prevalence*Mouse Abundance + Burn History Type |
| Prevalence ~ Adult Male Prevalence*Mouse Abundance + Year |
| Prevalence ~ Adult Male Prevalence*Mouse Abundance + PC1 |

Table S6. 28 *a priori* model candidates predicting infection viral load though a gamma hurdle model; where binomial model predicts the probability of being infected and the subsequent gamma model predicts the strength of infection, via ΔCT values.

| **Structure** |
| --- |
| Viral Load ~ 1 |
| Viral Load ~ Sex |
| Viral Load ~ Burn History Type |
| Viral Load ~ Condition |
| Viral Load ~ # Wounds |
| Viral Load ~ Year |
| Viral Load ~ Sex + Condition + Year +Burn History Type + # Wounds |
| Viral Load ~ Sex + Condition + Year + Burn History Type |
| Viral Load ~ Sex + Condition + Year |
| Viral Load ~ Sex + Condition |
| Viral Load ~ Sex + Year |
| Viral Load ~ Sex + Burn History Type |
| Viral Load ~ Sex + # Wounds |
| Viral Load ~ Burn History Type + Year |
| Viral Load ~ Sex*Condition |
| Viral Load ~ Sex*#Wounds |
| Viral Load ~ Sex*Condition +Burn History Type + # Wounds + Year |
| Viral Load ~ Sex*Condition + Year |
| Viral Load ~ Sex* Condition + Burn History Type |
| Viral Load ~ Sex*Condition + # Wounds |
| Viral Load ~ Sex*#Wounds + Burn History Type + Condition + Year |
| Viral Load ~ Sex*#Wounds + Burn History Type |
| Viral Load ~ Sex*#Wounds + Condition |
| Viral Load ~ Sex*#Wounds + Year |
| Viral Load ~ Year*Burn History Type |
| Viral Load ~ Year*Burn History Type + Sex |
| Viral Load ~ # Wounds + Condition |
| Viral Load ~ #Wounds*Condition |

Table S7. *A priori* model candidates for hurdle model predicting degree of wounding. Each structure was used to inform the hurdle model in all three stages: (i) predicting only the binomial step of the model (ii) predicting only the count step of the model (iii) predicting both the binomial and count steps.

| **Structure** |
|---|
| Wounding ~ 1 |
| Wounding ~ Sex |
| Wounding ~ SNV Infection |
| Wounding ~ Year |
| Wounding ~ Burn History Type |
| Wounding ~ Condition |
| Wounding ~ Viral Load |
| Wounding ~ Reproductive Status |
| Wounding ~ Sex + Condition |
| Wounding ~ Sex + SNV Infection |
| Wounding ~ Sex + Year |
| Wounding ~ Sex + Burn History Type |
| Wounding ~ Sex + Viral Load |
| Wounding ~ Condition + Burn History Type |
| Wounding ~ Condition + Reproductive Status |
| Wounding ~ Sex * Condition |
| Wounding ~ Sex + SNV Infection |
| Wounding ~ Sex * Reproductive Status |
| Wounding ~ Condition + Viral Load |
| Wounding ~ SNV Infection + Reproductive Status |

Table S8. Model outputs from GLM assessing mouse abundance

| Structure | K[1] | AICc[2] | ΔAICc[3] | ModelLik[4] | AICcWt[5] | LL[6] | Cum.Wt[7] |
|---|---|---|---|---|---|---|---|
| Relative Abundance[8] ~ Year[9] + PC1[10] | 4 | -61.120 | 0 | 1 | 0.373 | 36.782 | 0.373 |
| Relative Abundance ~ Proportion Reproductive[11] + PC1 | 4 | -60.600 | 0.521 | 0.771 | 0.287 | 36.522 | 0.660 |
| Relative Abundance ~ PC1 | 3 | -59.511 | 1.609 | 0.447 | 0.167 | 33.955 | 0.827 |
| Relative Abundance ~ Burn History Type[12] | 4 | -57.995 | 3.126 | 0.210 | 0.078 | 35.220 | 0.905 |
| Relative Abundance ~ Burn History Type + PC1 | 5 | -55.683 | 5.438 | 0.066 | 0.025 | 36.591 | 0.930 |
| Relative Abundance ~ 1 | 2 | -55.272 | 5.849 | 0.054 | 0.020 | 30.181 | 0.950 |
| Relative Abundance ~ Burn History Type + Proportion Reproductive | 5 | -55.054 | 6.067 | 0.048 | 0.018 | 36.277 | 0.968 |
| Relative Abundance ~ Burn History Type + Year | 5 | -52.963 | 8.158 | 0.017 | 0.006 | 35.231 | 0.974 |
| Relative Abundance ~ Burn History Type + Year + PC1 | 6 | -52.784 | 8.336 | 0.015 | 0.006 | 38.392 | 0.980 |
| Relative Abundance ~ Proportion Reproductive | 3 | -52.429 | 8.692 | 0.013 | 0.005 | 30.414 | 0.985 |
| Relative Abundance ~ Year | 3 | -52.345 | 8.775 | 0.012 | 0.005 | 30.373 | 0.990 |
| Relative Abundance ~ Burn History Type*Proportion Reproductive + PC1 | 8 | -51.870 | 9.250 | 0.010 | 0.004 | 48.335 | 0.993 |
| Relative Abundance ~ Burn History Type*PC1 + Proportion Reproductive | 8 | -51.086 | 10.034 | 0.007 | 0.002 | 47.943 | 0.996 |
| Relative Abundance ~ Burn History Type*Proportion Reproductive | 7 | -50.287 | 10.833 | 0.004 | 0.002 | 41.477 | 0.997 |
| Relative Abundance ~ Burn History Type*PC1 | 7 | -49.509 | 11.611 | 0.003 | 0.001 | 41.088 | 0.998 |
| Relative Abundance ~ Burn History Type + Year + PC1 + Proportion Reproductive | 7 | -48.753 | 12.368 | 0.002 | 0.001 | 40.710 | 0.999 |
| Relative Abundance ~ Proportion Reproductive + Year | 4 | -48.540 | 12.580 | 0.002 | 0.001 | 30.492 | 1.000 |
| Relative Abundance ~ Burn History Type*PC1 + Year | 8 | -44.814 | 16.306 | 2.88E-04 | 1.07E-04 | 44.807 | 1.000 |
| Relative Abundance ~ Burn History Type*PC1 + Proportion Reproductive + Year | 9 | -41.013 | 20.107 | 4.30E-05 | 1.60E-05 | 52.006 | 1.000 |
| Relative Abundance ~ Burn History Type*Proportion + Year | 8 | -39.217 | 21.903 | 1.75E-05 | 6.54E-06 | 42.009 | 1 |

Notes: [1]Number of model parameters [2]Akaike Information Criterion corrected for small sample sizes [3]Relative difference between best model and each other model in set [4]Relative likelihood of model given (exp(-0.5*delta[i])) [5]Akaike weights indicating the level of support of any given model being most parsimonious [6]log-likelihood of the model [7]Cumulative Akaike weights [8] Mouse relative abundance at sites [9]Year: 2023 or 2024 [10] PC1 loading from PCA analysis (Table

1) [11]Proportion of individuals that were identified as being sexually reproductive at time of capture [12] Burned, Unburned, or Reseeded

Abbreviations: PC1 = first principal component from habitat PCA

Table S9. Model outputs from GLM of individual infection status

| Structure | K[1] | AICc[2] | ΔAICc[3] | ModelLik[4] | AICcWt[5] | LL[6] | Cum.Wt[7] |
|---|---|---|---|---|---|---|---|
| SNV[8] ~ Burn History Type[9] | 3 | 248.804 | 0.000 | 1.000 | 0.242 | -121.362 | 0.242 |
| SNV ~ Burn History Type + Year[10] | 4 | 249.407 | 0.602 | 0.740 | 0.179 | -120.636 | 0.422 |
| SNV ~ Sex[11] + Burn History Type | 5 | 249.717 | 0.913 | 0.634 | 0.154 | -119.758 | 0.575 |
| SNV ~ Year* Burn History Type + Sex | 6 | 250.755 | 1.951 | 0.377 | 0.091 | -119.236 | 0.666 |
| SNV ~ Sex + Condition[12] + Year + Burn History Type | 6 | 250.788 | 1.984 | 0.371 | 0.090 | -119.252 | 0.756 |
| SNV ~ Year*Burn History Type | 5 | 251.338 | 2.533 | 0.282 | 0.068 | -120.568 | 0.825 |
| SNV ~ Sex*Condition + Burn History Type | 6 | 251.617 | 2.813 | 0.245 | 0.059 | -119.667 | 0.884 |
| SNV ~ Sex*#Wounds + Burn History Type | 6 | 251.635 | 2.831 | 0.243 | 0.059 | -119.676 | 0.943 |
| SNV ~ Sex + Condition + Year + Burn History Type + # Wounds[13] | 7 | 252.851 | 4.047 | 0.132 | 0.032 | -119.236 | 0.975 |
| SNV ~ Sex*#Wounds + Burn History Type + Condition + Year | 8 | 254.648 | 5.843 | 0.054 | 0.013 | -119.079 | 0.988 |
| SNV ~ Sex*Condition +Burn History Type + # Wounds + Year | 8 | 254.790 | 5.986 | 0.050 | 0.012 | -119.150 | 1.000 |
| SNV ~ Year | 2 | 269.357 | 20.552 | 3.44E-05 | 8.34E-06 | -132.658 | 1.000 |
| SNV ~ Sex + Year | 3 | 270.818 | 22.013 | 1.66E-05 | 4.02E-06 | -132.369 | 1.000 |
| SNV ~ Sex + Condition + Year | 4 | 272.822 | 24.017 | 6.09E-06 | 1.48E-06 | -132.344 | 1.000 |
| SNV ~ Sex*Condition + Year | 5 | 274.372 | 25.567 | 2.81E-06 | 6.80E-07 | -132.085 | 1.000 |
| SNV ~ Sex*#Wounds + Year | 5 | 274.711 | 25.906 | 2.37E-06 | 5.74E-07 | -132.254 | 1.000 |
| SNV ~ 1 | 1 | 286.034 | 37.229 | 8.24E-09 | 2.00E-09 | -142.010 | 1.000 |
| SNV ~ Sex | 2 | 287.048 | 38.243 | 4.96E-09 | 1.20E-09 | -141.504 | 1.000 |
| SNV ~ # Wounds | 2 | 287.845 | 39.041 | 3.33E-09 | 8.07E-10 | -141.903 | 1.000 |
| SNV ~ Condition | 2 | 287.846 | 39.041 | 3.33E-09 | 8.06E-10 | -141.903 | 1.000 |
| SNV ~ Sex + Condition | 3 | 288.961 | 40.156 | 1.91E-09 | 4.62E-10 | -141.440 | 1.000 |
| SNV ~ Sex + # Wounds | 3 | 289.017 | 40.213 | 1.85E-09 | 4.49E-10 | -141.468 | 1.000 |
| SNV ~ Sex*Condition | 4 | 290.101 | 41.297 | 1.08E-09 | 2.61E-10 | -140.983 | 1.000 |
| SNV ~ Sex*#Wounds | 4 | 291.045 | 42.241 | 6.72E-10 | 1.63E-10 | -141.455 | 1.000 |
| SNV ~ Sex*Condition + # Wounds | 5 | 292.070 | 43.265 | 4.03E-10 | 9.76E-11 | -140.934 | 1.000 |
| SNV ~ Sex*#Wounds + Condition | 5 | 292.975 | 44.171 | 2.56E-10 | 6.20E-11 | -141.387 | 1.000 |

Notes: [1]Number of model parameters [2]Akaike Information Criterion corrected for small sample sizes [3]Relative difference between best model and each other model in set [4]Relative likelihood of model given (exp(-0.5*delta[i])) [5]Akaike weights indicating the level of support of any given model being most parsimonious [6]log-likelihood of the model [7]Cumulative Akaike weights [8] SNV prevalence among individuals [9] Burned, Unburned or Reseeded [10]Year: 2023 or 2024 [11] Male or female [12] Size corrected mass, assessed through residuals of a linear regression of log-transformed hindfoot length and log-transformed weight [13] Number of visible wounds on each individual during capture

Table S10. Model output from GLM of site-level prevalence.

| Structure | K[1] | AICc[2] | ΔAICc[3] | ModelLik[4] | AICcWt[5] | LL[6] | Cum.Wt[7] |
|---|---|---|---|---|---|---|---|
| Prevalence[8] ~ Burn History Type[9] | 4 | -24.440 | 0.000 | 1.000 | 0.618 | 18.442 | 0.618 |
| Prevalence ~ Burn History Type + PC1[10] | 5 | -20.430 | 4.010 | 0.135 | 0.083 | 18.965 | 0.702 |
| Prevalence ~ 1 | 2 | -19.696 | 4.744 | 0.093 | 0.058 | 12.393 | 0.759 |
| Prevalence ~ Burn History Type + Year[11] | 5 | -19.571 | 4.869 | 0.088 | 0.054 | 18.536 | 0.813 |
| Prevalence ~ Burn History Type + Relative Abundance[12] | 5 | -19.457 | 4.983 | 0.083 | 0.051 | 18.479 | 0.865 |
| Prevalence ~ Relative Abundance | 3 | -18.640 | 5.800 | 0.055 | 0.034 | 13.520 | 0.899 |
| Prevalence ~ Proportion Adult Male[13] | 3 | -17.870 | 6.570 | 0.037 | 0.023 | 13.135 | 0.922 |
| Prevalence ~ Year | 3 | -17.714 | 6.726 | 0.035 | 0.021 | 13.057 | 0.943 |
| Prevalence ~ Relative Abundance + Proportion Adult Male | 4 | -17.618 | 6.822 | 0.033 | 0.020 | 15.031 | 0.964 |
| Prevalence ~ PC1 | 3 | -16.391 | 8.049 | 0.018 | 0.011 | 12.395 | 0.975 |
| Prevalence ~ Proportion Adult Male*Relative Abundance | 5 | -15.849 | 8.591 | 0.014 | 0.008 | 16.675 | 0.983 |
| Prevalence ~ PC1 + Relative Abundance | 4 | -15.561 | 8.879 | 0.012 | 0.007 | 14.003 | 0.990 |
| Prevalence ~ PC1 + Proportion Adult Male | 4 | -13.826 | 10.614 | 0.005 | 0.003 | 13.135 | 0.993 |
| Prevalence ~ Relative Abundance + Proportion Adult Male + Year | 5 | -13.683 | 10.758 | 0.005 | 0.003 | 15.591 | 0.996 |
| Prevalence ~ Burn History Type + Relative Abundance + Year | 6 | -13.150 | 11.290 | 0.004 | 0.002 | 18.575 | 0.998 |
| Prevalence ~ Proportion Adult Male*Relative Abundance + PC1 | 6 | -10.286 | 14.154 | 8.44E-04 | 5.22E-04 | 17.143 | 0.999 |
| Prevalence ~ Proportion Adult Male*Relative Abundance + Year | 6 | -9.644 | 14.796 | 6.12E-04 | 3.79E-04 | 16.822 | 0.999 |
| Prevalence ~ Burn History Type*Relative Abundance | 7 | -8.312 | 16.128 | 3.15E-04 | 1.95E-04 | 20.489 | 1.000 |
| Prevalence ~ PC1 + Relative Abundance + Proportion Adult Male + Year | 6 | -7.759 | 16.681 | 2.39E-04 | 1.48E-04 | 15.879 | 1.000 |
| Prevalence ~ Burn History Type* Proportion Adult Male | 7 | -7.319 | 17.121 | 1.92E-04 | 1.18E-04 | 19.993 | 1.000 |
| Prevalence ~ Burn History Type*PC1 | 7 | -7.302 | 17.138 | 1.90E-04 | 1.17E-04 | 19.984 | 1.000 |
| Prevalence ~ Proportion Adult Male*Relative Abundance + Burn History Type | 7 | -5.626 | 18.814 | 8.21E-05 | 5.08E-05 | 19.146 | 1.000 |
| Prevalence ~ Burn History Type + Relative Abundance + Year + Proportion Adult Male | 7 | -4.519 | 19.921 | 4.72E-05 | 2.92E-05 | 18.593 | 1.000 |
| Prevalence ~ Proportion Adult Male*Relative Abundance + Year + PC1 | 7 | -1.625 | 22.815 | 1.11E-05 | 6.87E-06 | 17.146 | 1.000 |
| Prevalence ~ Burn History Type + Relative Abundance + Year + Proportion Adult Male + PC1 | 8 | 6.680 | 31.120 | 1.75E-07 | 1.08E-07 | 19.060 | 1 |

| | | | | | | | |
|---|---|---|---|---|---|---|---|
| Prevalence ~ Burn History Type*PC1 + Relative Abundance + Proportion Adult Male | 9 | 19.690 | 44.130 | 2.61E-10 | 1.62E-10 | 21.655 | 1 |
| Prevalence ~ Burn History Type*PC1 + Relative Abundance + Year | 9 | 20.361 | 44.801 | 1.87E-10 | 1.16E-10 | 21.319 | 1 |
| Prevalence ~ Proportion Adult Male*Relative Abundance + Year + PC1 + Burn History Type | 9 | 23.792 | 48.232 | 3.36E-11 | 2.08E-11 | 19.604 | 1 |
| Prevalence ~ Burn History Type*PC1 + Relative Abundance*Proportion Adult Male | 10 | 49.355 | 73.795 | 9.45E-17 | 5.85E-17 | 21.989 | 1 |
| Prevalence ~ Burn History Type*PC1 + Relative Abundance + Year + Proportion Adult Male | 10 | 50.024 | 74.464 | 6.77E-17 | 4.18E-17 | 21.655 | 1 |
| Prevalence ~ Burn History Type*PC1 + Year + Proportion Adult Male | 10 | 50.024 | 74.464 | 6.77E-17 | 4.18E-17 | 21.655 | 1 |

Notes: [1]Number of model parameters [2]Akaike Information Criterion corrected for small sample sizes [3]Relative difference between best model and each other model in set [4]Relative likelihood of model given (exp(-0.5*delta[i])) [5]Akaike weights indicating the level of support of any given model being most parsimonious [6]log-likelihood of the model [7]Cumulative Akaike weights [8] SNV prevalence among sites [9]Burned, Unburned, or Reseeded [10]PC1 loading from PCA analysis (see Table 1) [11]Year: 2023 or 2024 [12]Relative mouse abundance at each site [13]Proportion of adult male mice at a site relative total adult mice at site

Table S11. Model outputs of gamma hurdle model to predict viral load.

| Structure | K[1] | AICc[2] | ΔAICc[3] | ModelLik[4] | AICcWt[5] | LL[6] | Cum.Wt[7] |
|---|---|---|---|---|---|---|---|
| Viral Load ~ 1 | 3 | 385.518 | 0.000 | 1.000 | 0.815 | -189.523 | 0.815 |
| Viral Load ~ # Wounds | 6 | 391.475 | 5.958 | 0.051 | 0.041 | -188.863 | 0.856 |
| Viral Load ~ Year | 6 | 392.205 | 6.687 | 0.035 | 0.029 | -189.227 | 0.885 |
| Viral Load ~ Condition | 6 | 392.433 | 6.915 | 0.032 | 0.026 | -189.341 | 0.910 |
| Viral Load ~ Sex | 6 | 392.721 | 7.203 | 0.027 | 0.022 | -189.485 | 0.933 |
| Viral Load ~ Sex + # Wounds | 7 | 394.108 | 8.590 | 0.014 | 0.011 | -188.862 | 0.944 |
| Viral Load ~ Burn History Type | 7 | 394.333 | 8.816 | 0.012 | 0.010 | -188.975 | 0.954 |
| Viral Load ~ Sex + Year | 7 | 394.655 | 9.137 | 0.010 | 0.008 | -189.136 | 0.962 |
| Viral Load ~ Sex + Condition | 7 | 395.055 | 9.538 | 0.008 | 0.007 | -189.336 | 0.969 |
| Viral Load ~ #Wounds*Condition | 8 | 395.550 | 10.032 | 0.007 | 0.005 | -188.210 | 0.974 |
| Viral Load ~ Year*Burn History Type | 9 | 395.873 | 10.355 | 0.006 | 0.005 | -186.936 | 0.979 |
| Viral Load ~ Burn History Type + Year | 8 | 396.092 | 10.575 | 0.005 | 0.004 | -188.481 | 0.983 |
| Viral Load ~ Sex*#Wounds | 8 | 396.786 | 11.268 | 0.004 | 0.003 | -188.828 | 0.986 |
| Viral Load ~ Sex + Burn History Type | 8 | 396.939 | 11.422 | 0.003 | 0.003 | -188.904 | 0.989 |
| Viral Load ~ Sex + Condition + Year | 8 | 397.197 | 11.680 | 0.003 | 0.002 | -189.033 | 0.991 |
| Viral Load ~ Sex*Condition | 8 | 397.225 | 11.708 | 0.003 | 0.002 | -189.047 | 0.994 |
| Viral Load ~ Sex*Condition + # Wounds | 9 | 397.992 | 12.474 | 0.002 | 0.002 | -187.996 | 0.995 |
| Viral Load ~ Sex*#Wounds + Year | 9 | 398.568 | 13.051 | 0.001 | 0.001 | -188.284 | 0.996 |
| Viral Load ~ Year*Burn History Type + Sex | 10 | 398.829 | 13.312 | 0.001 | 0.001 | -186.915 | 0.997 |
| Viral Load ~ Sex*#Wounds + Condition | 9 | 399.183 | 13.665 | 0.001 | 0.001 | -188.591 | 0.998 |
| Viral Load ~ Sex*Condition + Year | 9 | 399.657 | 14.139 | 8.51E-04 | 6.93E-04 | -188.828 | 0.999 |
| Viral Load ~ Sex*Condition + Burn History Type | 10 | 401.540 | 16.023 | 3.32E-04 | 2.70E-04 | -188.270 | 0.999 |
| Viral Load ~ Sex*#Wounds + Burn History Type | 10 | 401.615 | 16.098 | 3.19E-04 | 2.60E-04 | -188.308 | 0.999 |
| Viral Load ~ Sex + Condition + Year + Burn History Type | 10 | 401.634 | 16.117 | 3.16E-04 | 2.58E-04 | -188.317 | 1.000 |
| Viral Load ~ Sex + Condition + Year +Burn History Type + # Wounds | 11 | 402.729 | 17.212 | 1.83E-04 | 1.49E-04 | -187.295 | 1.000 |
| Viral Load ~ Sex*Condition +Burn History Type + # Wounds + Year | 12 | 404.868 | 19.351 | 6.28E-05 | 5.12E-05 | -186.720 | 1.000 |
| Viral Load ~ # Wounds + Condition | 12 | 404.868 | 19.351 | 6.28E-05 | 5.12E-05 | -186.720 | 1.000 |

| | | | | | | | |
|---|---|---|---|---|---|---|---|
| Viral Load ~ Sex*#Wounds + Burn History Type + Condition + Year | 12 | 406.004 | 20.486 | 3.56E-05 | 2.90E-05 | -187.287 | 1 |

Notes: [1]Number of model parameters [2]Akaike Information Criterion corrected for small sample sizes [3]Relative difference between best model and each other model in set [4]Relative likelihood of model given (exp(-0.5*delta[i])) [5]Akaike weights indicating the level of support of any given model being most parsimonious [6]log-likelihood of the model [7]Cumulative Akaike weights [8]Viral load, measured by ΔCt values [9]Number of visible wounds on each individual during capture [10]Year: 2023 or 2024 [11]Size corrected mass, assessed through residuals of a linear regression of log-transformed hindfoot length and log-transformed weight [12]Male or female [13]Burned, Unburned or Reseeded

Table S12. Model outputs from truncated Poisson hurdle model predicting the presence and degree of wounding on deer mice for three model types: (i) *Presence only model (ii) **Degree of wounding only model (iii) ***Both presence and degree of wounding model.

| Structure | K[1] | AICc[2] | ΔAICc[3] | ModelLik[4] | AICcWt[5] | LL[6] | Cum.Wt[7] |
|---|---|---|---|---|---|---|---|
| Wounding[8] ~ Sex[9] + Viral Load[10]* | 4 | 110.016 | 0.000 | 1 | 0.293 | -50.608 | 0.293 |
| Wounding ~ Viral Load* | 3 | 110.677 | 0.661 | 0.719 | 0.211 | -52.103 | 0.504 |
| Wounding ~ Viral Load** | 3 | 111.228 | 1.212 | 0.546 | 0.160 | -52.379 | 0.663 |
| Wounding ~ Viral Load*** | 4 | 112.402 | 2.386 | 0.303 | 0.089 | -51.801 | 0.752 |
| Wounding ~ Condition[11] + Viral Load* | 4 | 112.562 | 2.546 | 0.280 | 0.082 | -51.881 | 0.834 |
| Wounding ~ Condition + Viral Load** | 4 | 113.082 | 3.065 | 0.216 | 0.063 | -52.141 | 0.898 |
| Wounding ~ Sex + Viral Load** | 4 | 113.388 | 3.372 | 0.185 | 0.054 | -52.294 | 0.952 |
| Wounding ~ Sex + Viral Load*** | 6 | 114.193 | 4.176 | 0.124 | 0.036 | -50.221 | 0.988 |
| Wounding ~ Condition + Viral Load*** | 6 | 116.432 | 6.415 | 0.040 | 0.012 | -51.341 | 1 |
| Wounding ~ Sex + SNV Infection[12]*** | 6 | 585.188 | 475.171 | 6.57E-104 | 1.93E-104 | -286.452 | 1 |
| Wounding ~ Sex*** | 4 | 585.355 | 475.338 | 6.05E-104 | 1.77E-104 | -288.610 | 1 |
| Wounding ~ Sex*Reproductive Status[13]* | 5 | 585.374 | 475.357 | 5.99E-104 | 1.75E-104 | -287.586 | 1 |
| Wounding ~ Sex*Reproductive Status*** | 8 | 585.399 | 475.383 | 5.91E-104 | 1.73E-104 | -284.455 | 1 |
| Wounding ~ Sex + Condition*** | 6 | 586.846 | 476.830 | 2.87E-104 | 8.41E-105 | -287.281 | 1 |
| Wounding ~ Reproductive Status* | 3 | 587.838 | 477.822 | 1.75E-104 | 5.12E-105 | -290.879 | 1 |
| Wounding ~ Sex + Burn History Type[14]*** | 8 | 587.934 | 477.918 | 1.67E-104 | 4.88E-105 | -285.722 | 1 |
| Wounding ~ Condition + Reproductive Status* | 4 | 588.304 | 478.288 | 1.38E-104 | 4.05E-105 | -290.085 | 1 |
| Wounding ~ SNV Infection + Reproductive Status* | 4 | 588.522 | 478.506 | 1.24E-104 | 3.64E-105 | -290.194 | 1 |
| Wounding ~ Sex + Condition* | 4 | 588.690 | 478.674 | 1.14E-104 | 3.34E-105 | -290.278 | 1 |
| Wounding ~ Sex* | 3 | 588.698 | 478.682 | 1.14E-104 | 3.33E-105 | -291.309 | 1 |
| Wounding ~ Sex + Burn History Type* | 5 | 589.024 | 479.008 | 9.65E-105 | 2.83E-105 | -289.411 | 1 |
| Wounding ~ Sex*SNV Infection*** | 8 | 589.027 | 479.011 | 9.64E-105 | 2.82E-105 | -286.269 | 1 |
| Wounding ~ Sex + SNV Infection* | 4 | 589.091 | 479.075 | 9.34E-105 | 2.74E-105 | -290.478 | 1 |
| Wounding ~ Sex + Year[15]*** | 6 | 589.302 | 479.285 | 8.40E-105 | 2.46E-105 | -288.509 | 1 |

| | | | | | | | |
|---|---|---|---|---|---|---|---|
| Wounding ~ Reproductive Status$^{***}$ | 4 | 589.735 | 479.719 | 6.77E-105 | 1.98E-105 | -290.800 | 1 |
| Wounding ~ SNV Infection + Reproductive Status$^{***}$ | 6 | 590.191 | 480.175 | 5.39E-105 | 1.58E-105 | -288.954 | 1 |
| Wounding ~ Sex*Condition$^{*}$ | 5 | 590.201 | 480.185 | 5.36E-105 | 1.57E-105 | -289.999 | 1 |
| Wounding ~ Sex*Condition$^{***}$ | 8 | 590.419 | 480.403 | 4.81E-105 | 1.41E-105 | -286.965 | 1 |
| Wounding ~ Sex + Year$^{*}$ | 4 | 590.710 | 480.694 | 4.16E-105 | 1.22E-105 | -291.288 | 1 |
| Wounding ~ Sex*SNV Infection$^{*}$ | 5 | 590.959 | 480.943 | 3.67E-105 | 1.07E-105 | -290.379 | 1 |
| Wounding ~ Sex + SNV Infection$^{**}$ | 4 | 591.241 | 481.224 | 3.19E-105 | 9.34E-106 | -291.553 | 1 |
| Wounding ~ Condition + Reproductive Status$^{***}$ | 6 | 591.420 | 481.404 | 2.91E-105 | 8.54E-106 | -289.568 | 1 |
| Wounding ~ Sex$^{**}$ | 3 | 591.842 | 481.826 | 2.36E-105 | 6.91E-106 | -292.881 | 1 |
| Wounding ~ Sex*SNV Infection$^{**}$ | 5 | 593.142 | 483.126 | 1.23E-105 | 3.61E-106 | -291.470 | 1 |
| Wounding ~ Sex + Condition$^{**}$ | 4 | 593.299 | 483.283 | 1.14E-105 | 3.34E-106 | -292.583 | 1 |
| Wounding ~ Sex + Year$^{**}$ | 4 | 593.736 | 483.720 | 9.15E-106 | 2.68E-106 | -292.801 | 1 |
| Wounding ~ Sex + Burn History Type$^{**}$ | 5 | 593.983 | 483.967 | 8.09E-106 | 2.37E-106 | -291.891 | 1 |
| Wounding ~ SNV Infection$^{***}$ | 4 | 595.045 | 485.029 | 4.76E-106 | 1.39E-106 | -293.455 | 1 |
| Wounding ~ SNV Infection$^{**}$ | 3 | 595.058 | 485.041 | 4.73E-106 | 1.38E-106 | -294.489 | 1 |
| Wounding ~ Sex*Reproductive Status$^{**}$ | 5 | 595.099 | 485.083 | 4.63E-106 | 1.36E-106 | -292.448 | 1 |
| Wounding ~ SNV Infection$^{*}$ | 3 | 595.173 | 485.157 | 4.46E-106 | 1.31E-106 | -294.546 | 1 |
| Wounding ~ 1$^{***}$ | 2 | 595.199 | 485.183 | 4.40E-106 | 1.29E-106 | -295.580 | 1 |
| Wounding ~ 1$^{**}$ | 2 | 595.199 | 485.183 | 4.40E-106 | 1.29E-106 | -295.580 | 1 |
| Wounding ~ 1$^{*}$ | 2 | 595.199 | 485.183 | 4.40E-106 | 1.29E-106 | -295.580 | 1 |
| Wounding ~ Sex*Condition$^{**}$ | 5 | 595.292 | 485.276 | 4.20E-106 | 1.23E-106 | -292.545 | 1 |
| Wounding ~ Burn History Type$^{*}$ | 4 | 595.987 | 485.970 | 2.97E-106 | 8.70E-107 | -293.926 | 1 |
| Wounding ~ Condition$^{*}$ | 3 | 596.060 | 486.044 | 2.86E-106 | 8.39E-107 | -294.990 | 1 |
| Wounding ~ Condition$^{**}$ | 3 | 596.408 | 486.392 | 2.41E-106 | 7.05E-107 | -295.164 | 1 |
| Wounding ~ SNV Infection + Reproductive Status$^{**}$ | 4 | 596.813 | 486.797 | 1.97E-106 | 5.76E-107 | -294.339 | 1 |
| Wounding ~ Condition + Burn History Type$^{*}$ | 5 | 596.897 | 486.880 | 1.88E-106 | 5.52E-107 | -293.347 | 1 |
| Wounding ~ Reproductive Status$^{**}$ | 3 | 597.082 | 487.066 | 1.72E-106 | 5.03E-107 | -295.501 | 1 |
| Wounding ~ Year$^{**}$ | 3 | 597.188 | 487.172 | 1.63E-106 | 4.77E-107 | -295.554 | 1 |

| | | | | | | | |
|---|---|---|---|---|---|---|---|
| Wounding ~ Year* | 3 | 597.239 | 487.223 | 1.59E-106 | 4.65E-107 | -295.580 | 1 |
| Wounding ~ Condition*** | 4 | 597.283 | 487.267 | 1.55E-106 | 4.55E-107 | -294.574 | 1 |
| Wounding ~ Burn History Type** | 4 | 597.388 | 487.372 | 1.47E-106 | 4.32E-107 | -294.627 | 1 |
| Wounding ~ Burn History Type*** | 6 | 598.231 | 488.214 | 9.67E-107 | 2.83E-107 | -292.973 | 1 |
| Wounding ~ Condition + Reproductive Status** | 4 | 598.260 | 488.244 | 9.53E-107 | 2.79E-107 | -295.063 | 1 |
| Wounding ~ Condition + Burn History Type** | 5 | 598.792 | 488.776 | 7.31E-107 | 2.14E-107 | -294.295 | 1 |
| Wounding ~ Year*** | 4 | 599.242 | 489.226 | 5.83E-107 | 1.71E-107 | -295.554 | 1 |
| Wounding ~ Condition + Burn History Type*** | 8 | 600.615 | 490.599 | 2.94E-107 | 8.60E-108 | -292.063 | 1 |

Notes: [1]Number of model parameters [2]Akaike Information Criterion corrected for small sample sizes [3]Relative difference between best model and each other model in set [4]Relative likelihood of model given (exp(-0.5*delta[i])) [5]Akaike weights indicating the level of support of any given model being most parsimonious [6]log-likelihood of the model [7]Cumulative Akaike weights [8]Presence and degree of wounding assessed through a truncated poisson hurdle model [9]Male or Female [10]Viral load, measured by ΔCt values [11]Size corrected mass, assessed through residuals of a linear regression of log-transformed hindfoot length and log-transformed weight [12]SNV positive or negative through RT-qPCR [13]Assessed as either reproductive or not reproductive [14]Burned, Unburned, or Reseeded [15]Year: 2023 or 2024 *Presence only model **Degree of wounding only model ***Both presence and degree of wounding model
